# Label-free optical detection of bioelectric potentials using electrochromic thin films

**DOI:** 10.1101/2020.05.16.099002

**Authors:** Felix S. Alfonso, Yuecheng Zhou, Erica Liu, Allister F. McGuire, Yang Yang, Husniye Kantarci, Dong Li, Eric Copenhaver, J. Bradley Zuchero, Holger Müller, Bianxiao Cui

**Affiliations:** Department of Chemistry, Stanford University, Stanford, CA 94305; Department of Neurosurgery, Stanford University, Stanford, CA 94305; Department of Physics, 366 LeConte Hall, University of California, Berkeley, CA 94720, USA; Molecular Biophysics and Integrated Bioimaging, Lawrence Berkeley National Laboratory, Berkeley, CA 94720, USA

## Abstract

Understanding how a network of interconnected neurons receives, stores, and processes information in the human brain is one of the outstanding scientific challenges of our time. The ability to reliably detect neuroelectric activities is essential to addressing this challenge. Optical recording using voltage-sensitive fluorescent probes has provided unprecedented flexibility for choosing regions of interest in recording neuronal activities. However, when recording at a high frame rate such as 500-1000 Hz, fluorescence-based voltage sensors often suffer from photobleaching and phototoxicity, which limit the recording duration. Here, we report a new approach, Electro-Chromic Optical REcording (ECORE), that achieves label-free optical recording of spontaneous neuroelectrical activities. ECORE utilizes the electrochromism of PEDOT:PSS thin films, whose optical absorption can be modulated by an applied voltage. Being based on optical reflection instead of fluorescence, ECORE offers the flexibility of an optical probe without suffering from photobleaching or phototoxicity. Using ECORE, we optically recorded spontaneous action potentials in cardiomyocytes, cultured hippocampal and dorsal root ganglion neurons, and brain slices. With minimal perturbation to cells, ECORE allows long-term optical recording over multiple days.

## Introduction

Reliable detection of neuroelectric activities has been instrumental in deciphering how neurons encode information, expanding our understanding of how electrical activities lead to physiological outcomes. Several categories of recording methods have been developed, each with its own advantages and limitations. With high sensitivity and fast temporal response, electrodes have been the gold standard for detecting neuroelectric activities. Intracellular recording, using the patch clamp or sharp electrode, affords large signals and recording from user-selected cells, is invasive and limited to recording only one or two cells. Extracellular recording using multielectrode arrays (MEAs) is non-invasive and compatible with parallel and long-term recording (1). The development of high-density MEA and CMOS technology has enabled parallel recording of thousands of locations or more (2). However, prefabricated electrode arrays are fixed in space and not flexible to record user-selected sites.

Optical detection of electrical activities provides the spatial flexibility of choosing the target cells or areas of interest. The GCaMP family of calcium sensors has been widely used to optically read out neuronal activities with excellent signal to noise ratio (3, 4). In the last decade, the development of voltage-sensitive fluorescent proteins (5, 6) and small potentiometric dyes (7, 8) has advanced optical recording. These voltage-sensitive fluorophores exhibit faster kinetics than the GCaMP calcium sensors and are thus able to record discrete action potentials. Voltage-sensitive proteins are also genetically encodable to allow recording from a specific cell population. Although their operational mechanisms vary, voltage-sensitive fluorophores rely on inserting optically active molecules into the cell membrane through genetic modification or chemical incorporation, which sometimes lead to membrane capacitance overloading (9–12). Compared with GCaMPs, voltage-sensitive fluorophores are usually less bright. When recording at a high frame rate such as 500-1000 frames/sec, fluorescence-based voltage sensors photobleach in seconds to minutes (13).

A label-free optical detection of neuroelectric activities would avoid limitations associated with fluorescence-based molecular probes while still allowing spatial flexibility. A number of techniques are being explored in this area. Surface plasmon resonance (SPR) imaging has been shown to be able to detect mechanical motions of the neuron in response to current injection-triggered action potentials (14, 15), which was similarly detected by optical coherence tomography in Aplysia ganglia neurons (16, 17), by atomic-force microscopy in mammalian neurohypophysis (18), and by full-field interferometric imaging in spiking HEK cells (19). The fact that cellular mechanical motion can also be induced by other cellular mechanisms such as cell migration or mechanical contraction in cardiomyocytes makes these methods sensitive to artifacts (20). In a different approach, an elegant study using nitrogen-vacancy centers near the surface of a diamond has directly detected the magnetic fields caused by current injection-triggered action potentials in excised giant axons (21). However, the method currently requires averaging hundreds of events, and is thus not yet able to detect individual and spontaneous activities in a complex network of cells, or record single spikes. The ultimate goal of label-free optical recording, and eventually even imaging, of spontaneous cell electric activities in a complex neural network has yet to be achieved.

In this work, we report the development of Electro-Chromic Optical REcording (ECORE) for label-free optical detection of cellular action potentials. Electrochromic materials reversibly change their optical absorption spectrum in response to an externally applied voltage. These materials - such as metal-organic frameworks (22), metal oxides (23), and conductive polymers (24) - have been used in commercial applications including “smart windows” (25) for aircraft, glass facades for energy-efficient buildings, paper displays, and in optical tools (26). Here, we make use of this voltage-dependent absorption to achieve optical readout of electrical signals applied by cells. By building a sensitive optical detection setup, we have used ECORE to successfully achieve label-free optical detection of spontaneous neuronal action potentials in culture and in brain slices.

## Results and Discussion

### Optical detection of electric potentials with electrochromic PEDOT films

Our choice of electrochromic material is Poly(3,4-ethylenedioxythiophene) polystyrene sulfonate (PEDOT: PSS) (27), a widely-used conductive polymer that is chemically stable, easy to process, and biocompatible (28). Indeed, PEDOT:PSS has already been used as an electrode coating material in electrophysiology MEA recording (29). A cell that fires an action potential induces a local voltage fluctuation in the range of a hundred microvolts that can be detected by an extracellular electrode in its vicinity (**Fig. 1a, left**). If the cell is near a PEDOT:PSS thin film, we expect its action potential to induce small changes in the absorption of the film, which can be detected optically (**Fig. 1a, right**). For sensitive optical detection, we focus a laser beam onto the region of interest (ROI) through a prism-coupled total internal reflection configuration (**Fig.1b**). A balanced differential photodetector rejects laser intensity noise by comparing the reflection of the sample against a reference beam derived from the same laser. Detecting the total-internal reflection rather the transmission leaves the upper surface of the specimen accessible (e.g, for imaging). Because the probing light does not pass through the culture medium, this approach is insensitive to spurious signals such as floaters in the medium or air-liquid surface ripples induced by vibrations.

**Figure 1.**
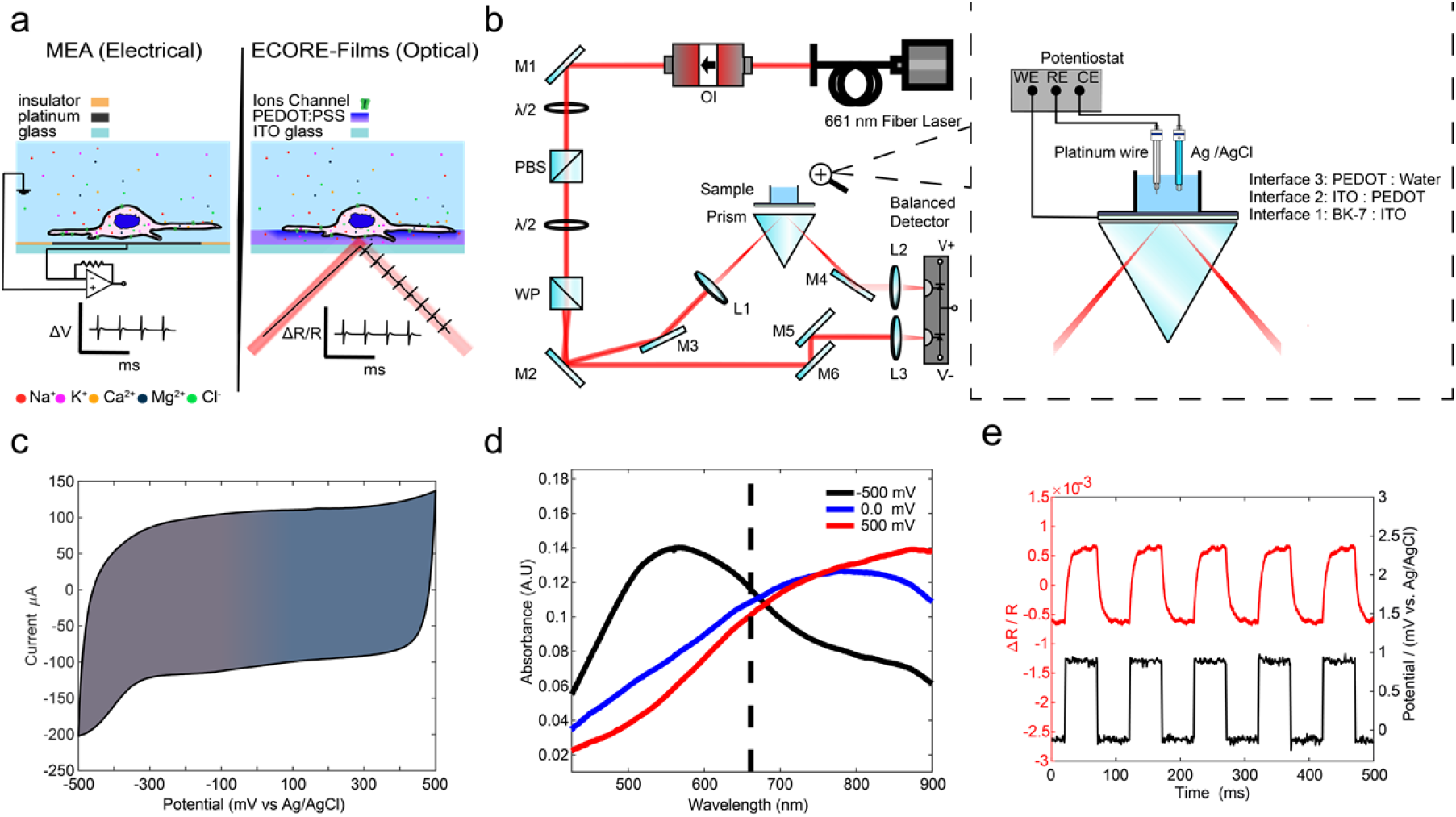
Characterization of the PEDOT:PSS film for ECORE optical recording. a) Schematics of MEA-recording and ECORE-recording from electrogenic cells. b) Optical set-up of ECORE with total internal reflection. The zoomed-in picture illustrates the interface and the electrical modulation of the electrochromic-film using a potentiostat. M: Mirror, PBS: polarizing beam splitter, WP: Wollaston Prism, L: lens, OI: optical isolator, and λ/2: half-wave plate. c) Cyclic voltammogram of PEDOT thin film in 0.1 M HEPES / KCl at a scan speed of 50 mV/s, d) UV-VIS absorption spectrum of the PEDOT:PSS film at three different applied voltages. e) The fractional reflectivity change (Δ*R*/*R, red curve*) is seen to be modulated by a 1-mV square wave applied to the film (*Potential, black curve*).

We produce PEDOT:PSS thin films (referred to as PEDOT hereafter) by electrodeposition directly from monomers onto an ITO-coated glass substrate (30). PEDOT films are chemically stable and have been shown to be stable in cell culture medium for months or longer (31, 32). We have observed that the film is stable for >3 months without any indication of decay. We characterized the electrochemical properties of PEDOT by cyclic voltammetry (33). The cyclic voltammogram shows no pronounced peaks indicative of faradaic current (**Fig. 1c**). Therefore, the PEDOT film resembles a pseudocapacitor electrochemically. The electrochromism of PEDOT arises from modulation of its π-conjugated system by an electric potential (34) (see supplementary text, **Fig. S1-2**). Absorption spectra of the film show a clear voltage-dependent shift in the absorption maximum. A −500 mV (all potentials are vs. Ag/AgCl electrode) results in a maximum absorption at λ_max_ = 580 nm, while a positive bias of +500 mV results in λ_max_ = 850 nm (**Fig. 1d**). Upon application of a train of 1 mV square waves to a 1 cm^2^ film, we detected a clear change in the reflection signal using a 660-nm probing laser. The fractional reflectivity change (Δ*R*/*R*) shows a train of optical responses that closely follow the train of applied electric voltages (**Fig. 1e**). Although the 660 nm wavelength does not lead to the highest sensitivity for voltage-changes, it is sufficient to detect small changes induced by cellular electrical signals as we show below.

### Characterization and optimization of ECORE

Detecting cellular electrical signals requires a sensitivity of 10 μV or better, 4-5 orders of magnitude less than the voltage (hundreds of mV) required to induce a color change of PEDOT:PSS that would be visible to the naked eye (35, 36). To maximize the relative reflectivity change, and thus the optical detection sensitivity, we built a four-layer (glass:ITO:PEDOT:water) model to study how the light reflectivity is modulated by changes in the extinction coefficient of the PEDOT film (See supplementary text, **Fig. S3-4**). We first optimized Δ*R*/*R* by tuning the incident angle. As the incident angle increases, the measured *ΔR/R* first increases and then decreases (**Fig. 2a**). The optimal incident angle is about 67.0°, which is larger than the critical angle of 62.5 and in good agreement with the model. We further optimize Δ*R*/*R* by precisely controlling the thickness of the PEDOT layer (37). The optical contrast first increases with the film thickness as more sensing material is added, but then starts to decrease roughly when the thickness exceeds the penetration of evanescent light at the interface. The optimal thickness to achieve the highest contrast is determined to be about 115 nm (**Fig. 2b**). By fixing the optimal incident angle and the film thickness, we measure the optical contrast *ΔR/R* as a function of bias voltage (*V*_0_) while maintaining Δ*V=*1 mV (Fig. 2c). The contrast shows a maximum at *V*_0_ = −100 mV, a minimum at *V*_0_ = +100 mV, and a good sensitivity *V*_0_= 0 mV. In the cell culture medium, the open-circuit potential of our system (−50 mV) brings the film closer to the optimum sensitivity without any active bias.

**Figure 2.**
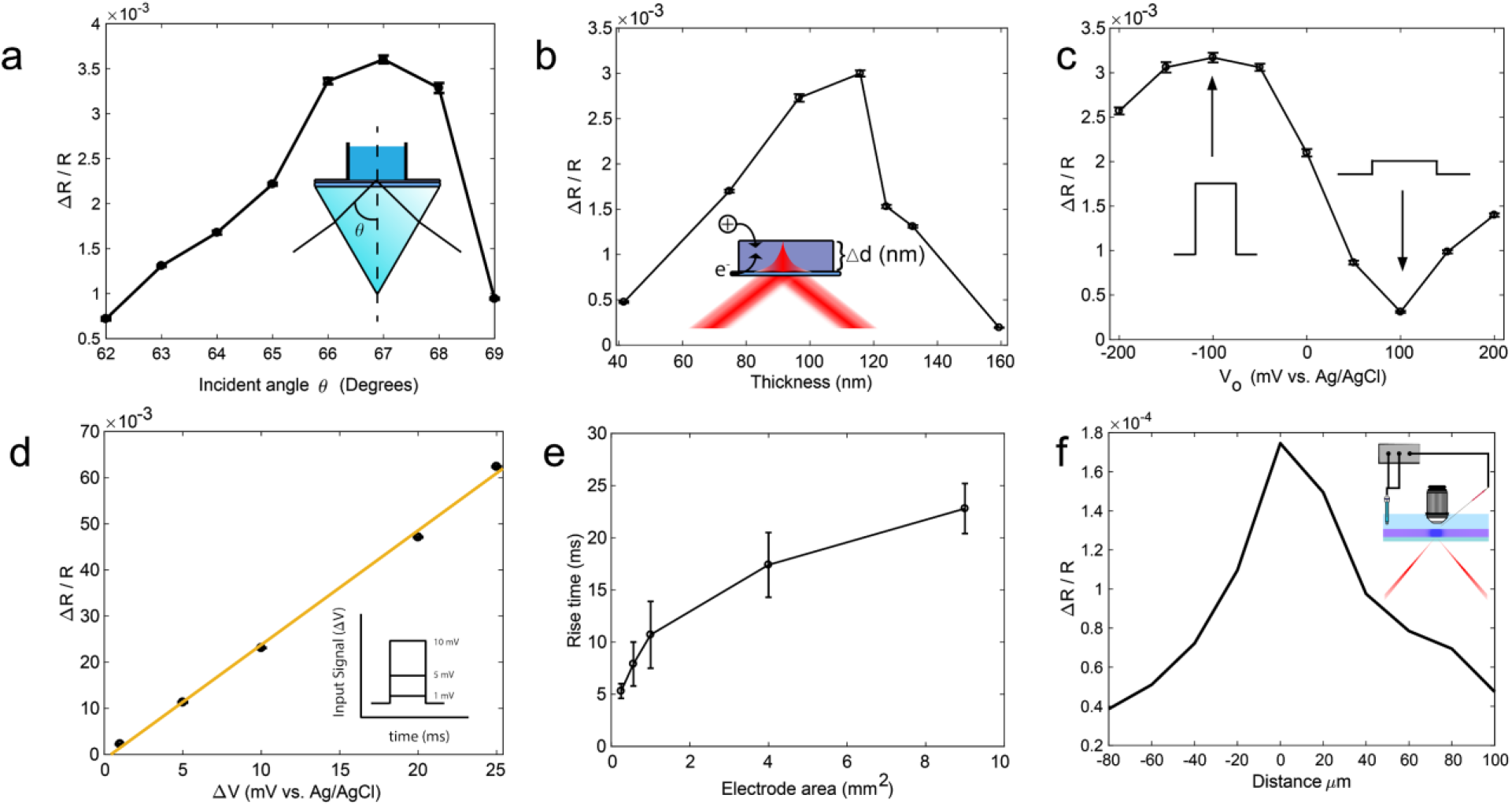
Optimization of ECORE. Change in fractional reflectivity change (Δ*R*/*R*) as a function of a) incident angle *θ* (*V*_0_= 0 and Δ*V* = 1 mV at 10 Hz), b) film thickness (*V*_0_= 0 and Δ*V* = 1 mV at 10 Hz), and c) bias voltage (*θ* = 67° and Δ*V* = 1 mV at 10 Hz). d) Δ*R*/*R* as a function of Δ*V* (*d* =120 nm, *V*_0_= 0 and *θ* = 67°). e) The film response time scales with the square root of the film’s surface area. f) The change in reflectivity (Δ*R*/*R*) decreases as the laser spot moves away from the microelectrode that locally applies 1-mV square wave voltage.

Finally, the optical contrast *ΔR/R* is linearly proportional to the applied voltage Δ*V* over a wide range at bias *V*_0_ = 0 (**Fig. 2d**). This linear dependence allows quantitative conversion of optical responses to voltages if the film is calibrated. For this work, we directly use *ΔR/R* instead of converting to voltages. Due to the instrument limit of the potentiostat, we are unable to apply electric pulses smaller than 1 mV. With a typical signal *ΔR/R* ~3.0 × 10^−3^ per 1 mV applied voltage and a typical standard deviation of the noise at 2.0 ×10^−5^ or less, we estimate a detection sensitivity of 6.7 μV with zero bias with a sampling rate of 10 kHz.

The temporal response of the film must be sufficiently fast in order to capture cellular electric potentials in the order of 1 ms. However, the long response time of macroscopic electrochromic coatings, typically on the order of seconds (38, 39), is too slow for capturing millisecond neuroelectric activities. We hypothesize that the temporal response is determined by charging of the active electrode area and the bulk diffusion of the counter ions necessary for charge stabilization (40). To test the hypothesis and establish a relationship between the film area and the temporal resolution, we measured the optical response time vs. PEDOT film areas. The response time is defined as the time for *ΔR/R* to rise from 10% to 90% of its final value after subjecting the film to a 1 mV square-wave at 0 mV bias potential. The PEDOT film area is precisely controlled from 9 mm^2^ to 0.25mm^2^ by patterning the ITO area through photolithography (see Supplementary information). Our measured temporal response of the film scales with the square-root of the PEDOT area *A^1/2^* (**Fig 2e**). This measurement agrees with an RC circuit model for microelectrodes (41), where τ=*RC*, where *R* is the electrolyte resistance proportional to *1/A^1/2^* and *C* the double-layer capacitance, proportional to *A*. We also experimentally measured the electrochemical impedance spectrum for each film and then modeled the impedance data using the equivalent circuit described above (**supplementary Fig. S5, Table S1**). The calculated response time from the impedance data agrees very well with our direct optical measurement. Therefore, the temporal resolution improves significantly when the active area becomes smaller. For a cell cultured on a PEDOT film, the active film area that is affected by cellular action potentials is similar to the cell size. We thus expect a response time of less than 0.2 ms for a cell area of 400 μm^2^.

The spatial resolution of the PEDOT:PSS film must be sufficient to detect individual cells. When a cell is in close contact with the film, the cell’s electrical activity will charge the film locally. The reported charge mobility in PEDOT:PSS (42, 43) indicates that the charge diffusion in 1 ms is about 10^−3^-10^−1^ μm^2^, much smaller than the size of a cell. We experimentally determine the spatial resolution by measuring optical responses at different film locations when an electrical pulse is applied locally through a microelectrode a few micrometers above the film surface (**Fig. 2f, supplementary information**). The optical response decays quickly as the laser spot moves away from the microelectrode. The measured full width at half maximum for the optical response is 33.4 μm, which is the size of the laser spot. Therefore, the spatial resolution of the EC film is limited not by the charge diffusion, but by the size of the probing laser spot. This size has been chosen to match the approximate size of the cell and can be further reduced if desired.

For all the cell studies, our ECORE measurements were carried out in an open circuit configuration (−50 mV open circuit potential) without any active electrode in the solution. Cells were cultured on 1 cm^2^ PEDOT films for a few days to weeks before ECORE measurements. We first used ECORE to detect action potentials in monolayers of human iPSC-derived cardiomyocytes (**Fig. 3a**). Using ECORE with a 10 kHz sampling frequency, we optically detect large and periodic optical signals (**Fig. 3b**). However, these large signals are due to the mechanical contraction of these cells that accompanies each action potential through excitation-contraction coupling. These mechanical signals arise when the movement of the cell membrane modifies the boundary conditions at the interface; they can be measured in the absence of the PEDOT layer. Careful examination of the ECORE trace reveals that, about 15 ms before the start of each mechanical contraction, there is a sharp spike resembling extracellularly-recorded action potentials (**arrows, Fig. 3c**). To confirm that these sharp spikes were indeed action potentials, we treated cardiomyocytes with 12.5μM blebbistatin (12.5μM) while the cells were being continuously recorded by ECORE (**Fig. 3d**). Blebbistatin is a potent myosin inhibitor that is known to inhibit cardiomyocyte contraction without eliminating its action potential (44). Magnified parts of the ECORE data after 0 min, 5 min, and 10 min of drug treatment show a drastic decrease of the mechanical signal at 5 min and then an almost complete elimination at 10 min (**Fig. 3e**). However, the sharp spikes corresponding to action potentials were unaffected by the blebbistatin treatment (**Fig. 3f,** magnified parts of the corresponding spikes in Fig. 3e). This confirms that ECORE is able to reliably detect individual action potentials in cardiomyocytes. The electrical and mechanical signals are clearly distinguished by their timing and shape.

**Figure 3.**
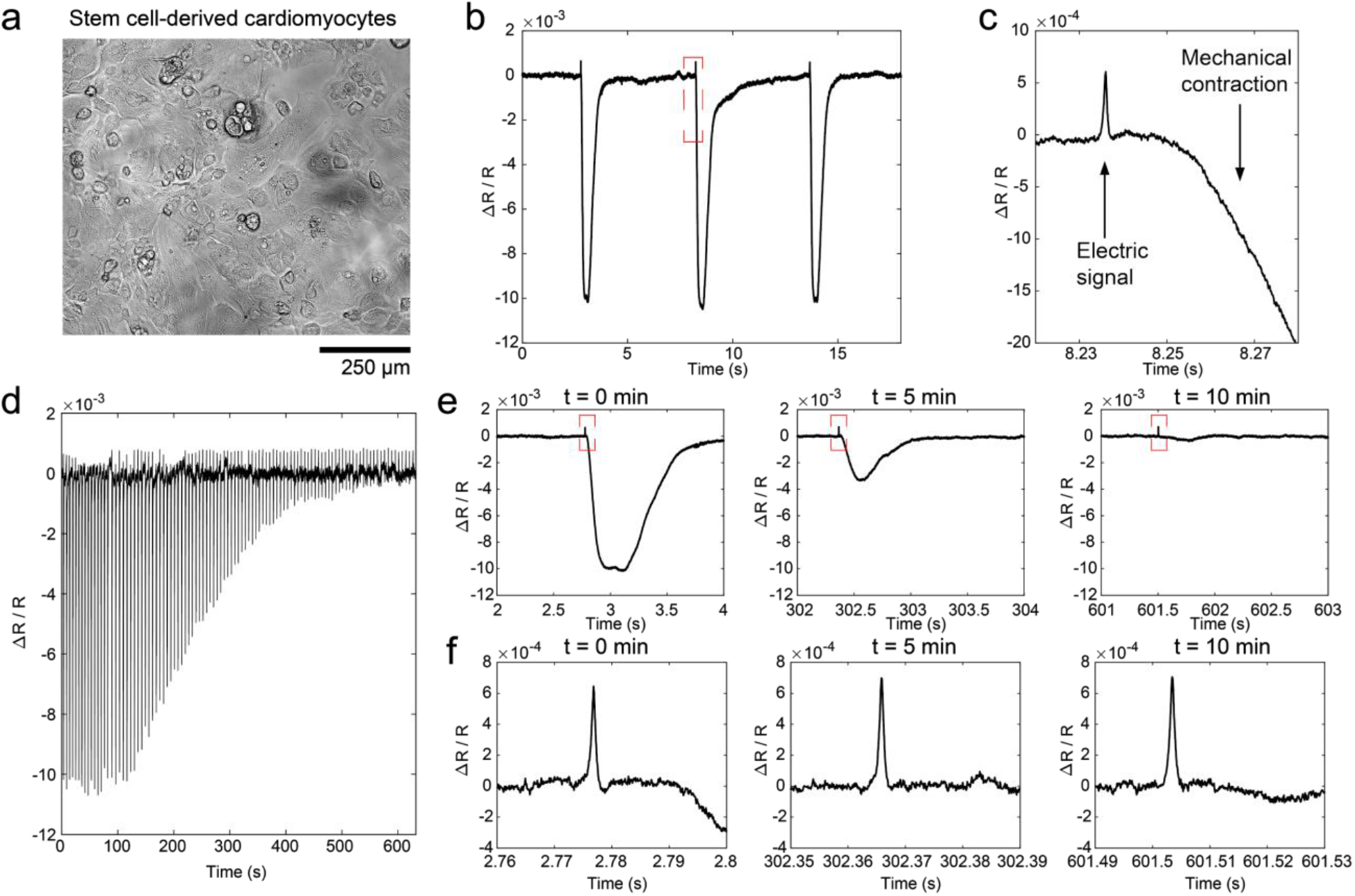
ECORE optical recording in stem cell-derived cardiomyocytes. **a**) Bright-field image of stem-cell derived cardiomyocytes cultured in a PEDOT film. **b**) Typical ECORE optical recording of cardiomyocytes. **c**) Enlarged part of b) shows the electrical signal as a small spike occurring about 15 ms before the onset of the cell’s mechanical contraction. **d**) ECORE recording shows a gradually decreased signal upon the application of 12.5 μM blebbistatin over 10 min. **e**) Enlarged parts of d) at 0 min, 5 min, and 10 min, respectively, show a drastic decrease of the mechanical contraction. **f**) Zoomed parts of (e) show that electric spikes remain the same at 0, 5, and 10 min.

We have used standard multi-electrode array (MEA) recording technique to confirm that our iPSC-derived cardiomyocytes exhibited spontaneous and periodical action potentials (**Fig. 4a**). MEA recording shows both monophasic spikes and biphasic electrical spikes at 10 kHz sampling frequency. As previously reported, the amplitude and waveform of action potential spikes strongly depends on the coupling and the relative location between the electrode and the cell (45). Using ECORE, we also detected both monophasic action potential signals (e.g. ROI 1 trace in **Fig. 4b**) and biphasic action potential signals (e.g. ROI 2 trace in **Fig. 4b**). Similar to MEA, the waveform and amplitude of ECORE depends on the coupling of the cells to the surface and the resulting ionic currents. The signal-to-noise ratio for ECORE recording (**Fig. 4b**) is similar to that of MEAs (**Fig. 4a**) at 10 kHz recording bandwidth and is better than fluorescence-based optical recording by voltage-sensitive proteins or dyes at the maximum 1kHz bandwidth. It is also orders of magnitude better than previous label-free optical recording methods and thus enables recording of single, spontaneous, and un-averaged action potentials.

**Figure 4.**
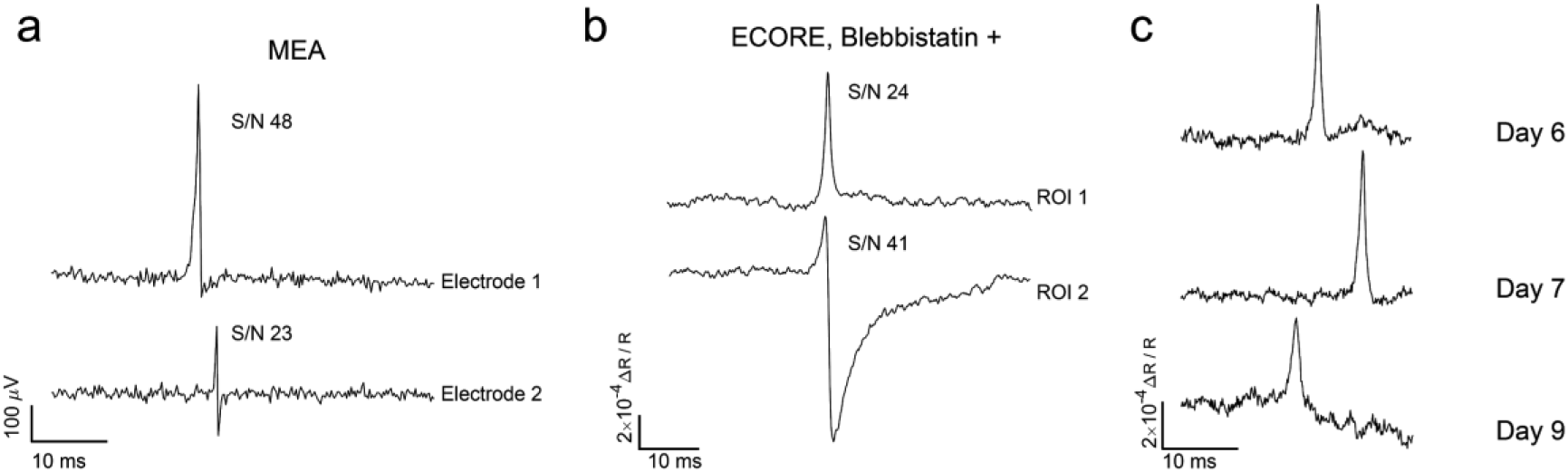
ECORE provides high S/N that is comparable to MEA recording. a) MEA electric recording shows both monophasic and biphasic signals with indicated S/N and b) ECORE optical recording also shows both monophasic and biphasic signals with an S/N that is comparable to MEA recording. c) ECORE allows multiple days *in vitro* (DIV) recordings of the same batch of cardiomyocytes cells.

ECORE is well-suited for long term recording of the electrical activities in cardiomyocytes since it requires no modification of the cell and does not suffer from photobleaching. With a 10 kHz recording bandwidth, we usually measure 10 min or more for each optical recording session. This duration, limited only by the lack of an appropriate culture chamber, is much longer than what can be achieved using voltage-sensitive fluorophores. Furthermore, we are able to measure the same culture for many days over multiple recording sessions. **Figure 4c** shows ECORE recordings from the same culture of cardiomyocytes at different locations at 6, 7 and 9 days on the PEDOT surface cultured *in vitro* (DIV).

After validating our platform with cardiomyocytes, we proceeded to record neuronal action potentials in cultured hippocampal neurons (**Fig. 5a**). Dissociated hippocampal neurons from embryonic E18 rats were cultured on the PEDOT film until they formed a network *in vitro* and exhibited spontaneous action potentials. ECORE measurements were made three weeks after plating when hippocampal neurons exhibit spontaneous and sparse electric activities (**Fig. 5b**). To confirm that the recorded signals in **Fig. 5b** were indeed neural electrical activity, we treated the same culture with carbachol, an activator of muscarinic acetylcholine receptors (mAChRs) that is known to increase the firing rate of hippocampal neurons (46). Upon 20 μM bath application of carbachol, the hippocampal neurons exhibited an increase in complex and unsynchronized electric signals (**Fig. 5c**), which agrees well with what has been reported in the literature (1). We note that the increase in the firing rate by application of carbachol was accompanied by a decrease in the amplitude of the electrical spikes which also agreed with previous literature report (47).

**Figure 5.**
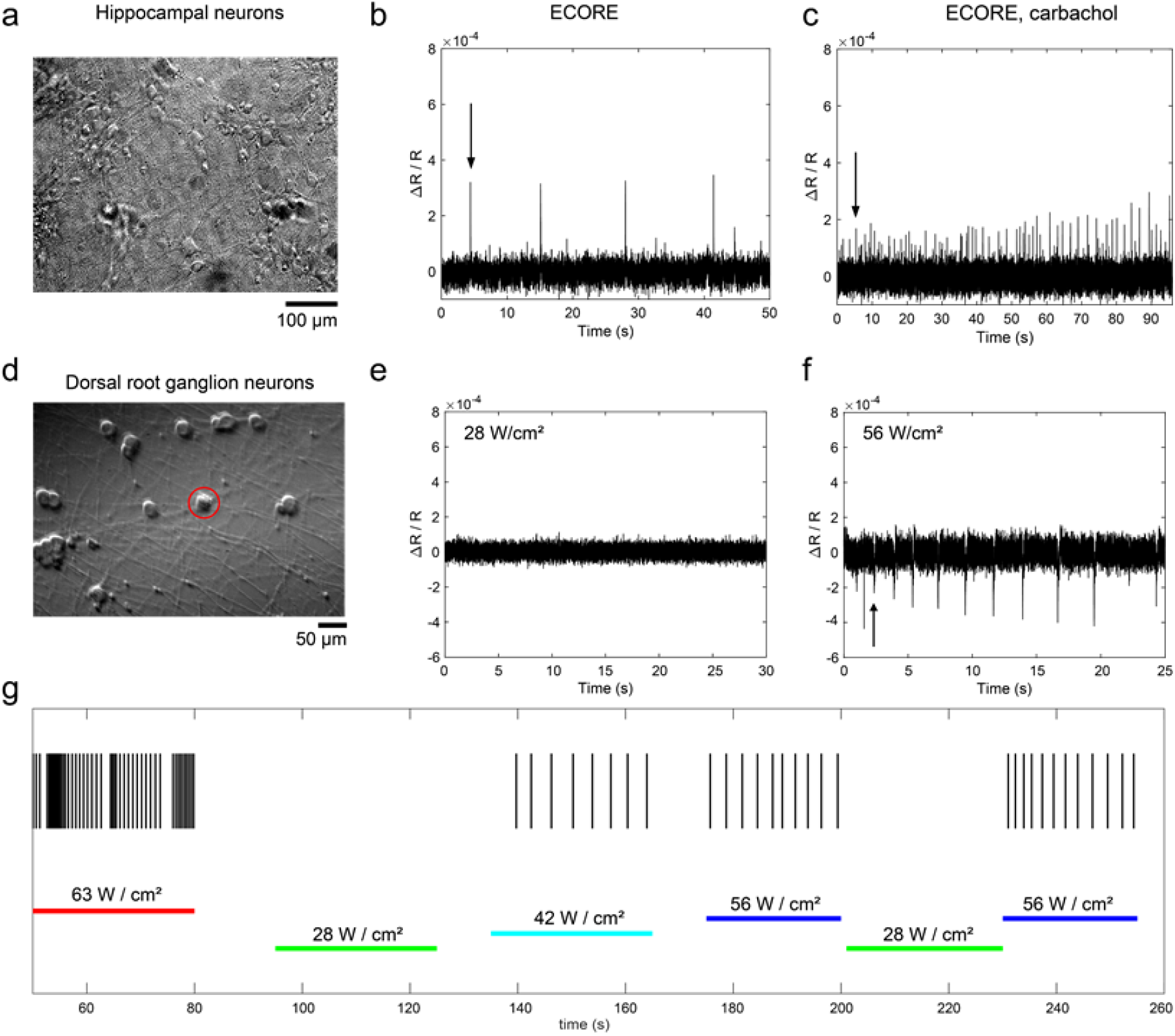
ECORE optical recording in cultured hippocampal neurons and dorsal root ganglion (DRG) neurons. **a)** Bright-field image of the hippocampal neurons cultured on a PEDOT film, **b)** ECORE optical recording of spontaneous activity in hippocampal neurons. **c)** At the same recording site in b), the application of 20 μM of carbachol significantly increased the firing frequency. **d)** Bright-field image of DRG neurons cultured on a PEDOT film. **e)** No spontaneous DRG activity was observed at the standard ECORE recording optical intensity of 28 W/cm^2^. **f)** Induced electric activity can be observed at a light intensity of 56 W/cm^2^. **g)** A continuous ECORE recording of the same location while the laser optical intensity was modulated from 63 to 28, 42, 56, 28, and 56 W/cm^2^, with each period lasting 25-30 s. Each vertical short line indicates an electric activity at the indicated time.

Dorsal root ganglion (DRG) neurons are sensory neurons known for the sensitivity to mechanical (48), chemical (49), thermal (50) and noxious stimuli. Embryonic rat DRG neurons were dissected, isolated, purified, and cultured according to the protocol established by the Zuchero group (51). We cultured DRG neurons on poly-D-lysine/laminin coated PEDOT film using the same protocol. After a week of culture, the brightfield image shows a cell body about 30 μm in diameter with long axons that are projecting out (**Fig. 5d**). Embryonic DRG neurons have been reported to exhibit low spontaneous activities (52, 53). Indeed, even with repeated effort, we were unable to detect spontaneous activity from these DRG neurons using the standard light intensity at 28 W/cm^2^ (**Fig. 5e**). However, when the optical intensity was doubled to 56 W/cm^2^, we observed an increase in the electric activities from these neurons (**Fig. 5f**). The firing pattern of each DRG neuron tested (n=21) differed between cells; however, we always observed an increase in the electric activities at high optical power. Many previous studies have demonstrated that light-absorbing materials can convert light to heat for photothermal stimulation of neurons (54–56). PEDOT absorbs 660 nm light strongly (57), thus we hypothesize that higher optical intensity might be locally heating up PEDOT and thus thermally activate the DRG neurons.

To confirm that electric activities of DRG neurons were indeed modulated by the laser optical intensity we increased the laser intensity to values well above the amount required for recording purposes. We sequentially changed the intensity from 63 to 28, 42, 56, 28, and 56 W/cm^2^ with each period lasting 25-30 s during a continuous 5-min ECORE session of the same DRG neuron (**Fig. 5g**, horizontal lines). The firing activity appears to be strongly dependent on the laser optical intensity (**Fig. 5g**, each vertical short line indicating an electric activity at the indicated time). No activity is detected during the two periods at 28 W/cm^2^ and mild activities are detected at the three periods with 42 and 56 W/cm^2^. Strong activities are detected at the period at 63 W/cm^2^. The photothermal stimulation on DRG neurons using intensities of up to 56 W/cm^2^ is fully reversible, except at the highest intensity of 63 W/cm^2^, where it is only reversible after a short duration such as shown in **Fig. 5g**. With longer illumination at 63 W/cm^2^, DRG neurons activities completely and irreversibly stopped, likely due to cell damage caused by over-stimulation. At moderate optical intensity (42-56 W/cm^2^), ECORE can be used to photothermally stimulate and electrochemically record cells without any genetic manipulation. A previous report (58) showed that photo illumination at 5900 W/cm^2^, λ = 640 nm for 240 s led to minimal phototoxicity in cultured mammalian U2OS cells.

For normal ECORE recording, we used an intensity of less than 28 W/cm^2^ at λ = 660 nm, well below the threshold that would cause direct cell damage or photostimulation. In a cell free system, we measured |Δ*R*/*R*| for 1-mV square waves as a function of the light intensity (**Fig. S6**). |Δ*R*/*R*| is relatively stable at low optical intensities but decreases as the optical intensity increases above 30 W/cm^2^. This decrease in |Δ*R*/*R*| is a signature that the PEDOT film is out of the linear absorption regime and leads to a reduced recording quality at higher optical power. Therefore, we avoid going to higher light intensities except for photothermal-simulation purposes.

The brain slice is a widely-used model in the field of molecular and cellular neuroscience (**Fig. 6a**). With their local circuits and the cyto-architecture relatively intact, they have been employed in pharmacological studies in controlled environments and electrophysiological studies to characterize properties of individual neurons and neuronal networks. We measured the electrical activity of hippocampal brain slices obtained from P7 rats and cultured using Stoppini’s method (59). After 10 days of culture on a semipermeable membrane, the brain slice is flipped onto either a PEDOT film or a MEA device. The tissue is gently pressed onto the surface using a 3D-printed adapter terminated with a nylon mesh and immersed in artificial cerebrospinal fluid (ACSF) saturated with carbogen gas. We first confirm that these brain slices exhibit spontaneous electric activities by using the conventional MEA device. MEA recording shows sparse and sporadic spikes (**Fig. 6b**), which is typical of hippocampus slice recording (60).

**Figure 6.**
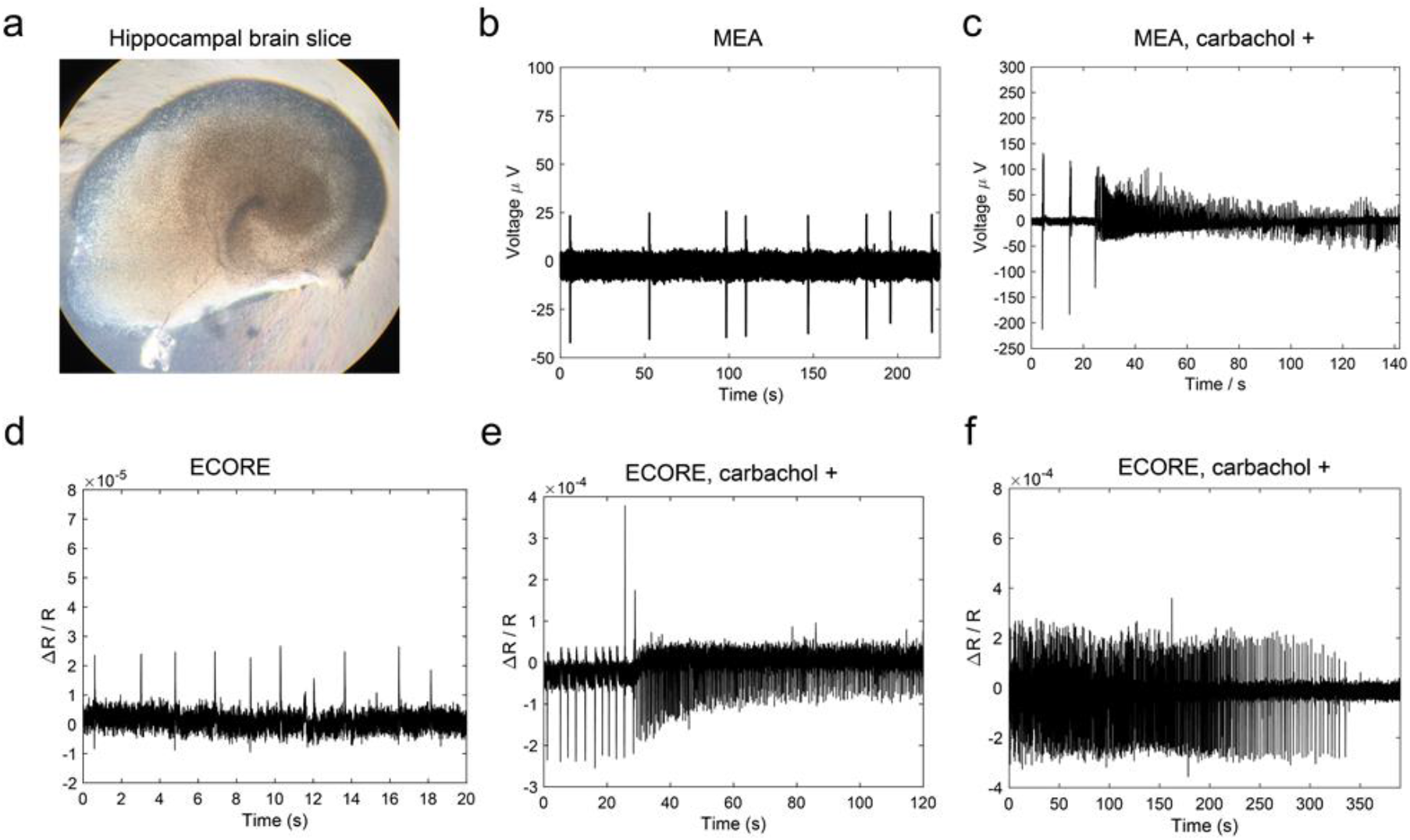
ECORE optical recording and MEA electric recording of hippocampal brain slices with insets showing representative electrical activities. a) Brightfield image of a cultured hippocampal brain slice. b) MEA electric recording shows sporadic spikes of neuronal activities from a brain slice. c) After a brain slice is exposed to 20 μM of carbachol, MEA recording shows a strong increase in the spiking frequency. d) ECORE optical recording shows sparse and spontaneous spiking activities from a brain slice. e) ECORE optical recording shows a drastic increase in spiking frequency after 20 μM of carbachol treatment. f) ECORE optical recording shows the carbachol-induced high spiking frequency gradually decreases during hypoxia.

After application of 20 μM carbachol, an agonist of the acetylcholine receptor, we observed a significant increase in the spike frequency by MEA recording (**Fig. 6c**), consistent with previous reports (61). Similar to the MEA recording, our ECORE recording shows sparse and spontaneous spiking patterns from brain slices (**Fig. 6d**). The spikes are sometimes monophasic and sometimes biphasic. The peak-to-peak amplitude of Δ*R*/*R*=3×10^−5^ can be observed with a S/N ratio of 14 at 1 kHz (−3dB) bandwidth. To confirm that the optically-recorded spikes are indeed from the electrical activity of the brain slice, we apply 20 μM carbachol to the brain tissue while it is being continuously recorded on the optical setup. We observe a sudden and pronounced increase in the optically-recorded spiking frequency (**Fig. 6e**), in agreement with our MEA measurements (**Fig. 6c**). Lastly, the electrical activity of brain slices depends on the availability of oxygen; hence hypoxic conditions decrease the electrical activity of brain slices. Indeed, we find that the optical signal of a brain slice exposed to 20 μM carbachol without a continuous supply of carbogenated ACSF gradually decreases and ceases (**Fig. 6f**). Re-saturating the ACSF with carbogen restores the electrical activity of the same brain slice (supplementary Fig. S7). These results demonstrate that we have successfully achieved label-free optical recording of hippocampal brain slices.

In this work, we develop a method that achieves label-free optical detection of bioelectric activities. Using electrochromism, a unique material property, ECORE provides the spatial flexibility of optical detection without requiring molecular insertions or genetic modifications to the cells. We note that ECORE is an extracellular method like MEAs, and thus is not genetically encodable and cannot be used to measure absolute voltages such as resting membrane potentials. Nevertheless, it offers complementary advantages to fluorescence-based voltage recording and would be valuable for measuring cells that are sensitive to molecular perturbation or difficult to modify genetically, such as human stem cell derived neurons for disease modeling. ECORE does not suffer from photobleaching or phototoxicity, and thus enables long recording sessions; e.g. we regularly record 20-minute or longer at 10kHz sampling rate and have recorded the same culture in multiple sessions over a week or longer.

This work demonstrates the feasibility to optically detect action potentials via electrochromic materials using a single point of detection. Extending this work to imaging can be achieved by utilizing an acousto-optic deflector (AOD) to steer the beam to scan multiple sites of interest for simultaneous detection of many cells. For example, two-dimensional AODs could scan 100 spots at a rate of 1 kHz. The sensitivity of ECORE can be further improved by choosing an optimal wavelength, polarization, and by using multiple probing wavelengths for noise rejection. The fast kinetics, long-term stability, and non-invasive nature makes ECORE an attractive candidate as a complementary tool applied toward the growing field of all-optical electrophysiology.

## Materials and Methods

Detailed method and supporting figures can be found in SI Materials and Methods.

## Author Contributions

F.A. designed and performed most experiments, interpreted data, and drafted the manuscript. Y.Z. and E.L. helped perform experiments and data collection. A.F.M. helped troubleshooting the apparatus. Y.Y. provided assistance in the electrical characterization of samples and establishing a cell culturing protocol. H.K. and J.B.Z. provided the DRG neurons for the optical recording. D.L. prepared the hippocampal brain slices. E.C. helped interpretation and assisted manuscript writing. H.M. built the theory and model, participated in building the optical setup and the writing of the manuscript. B.C. initiated the idea, participated in the experimental design, data interpretation, and manuscript writing.

## Author Information

All data needed to evaluate the conclusions in the paper are present in this paper and the supplementary materials. The authors declare no competing interests.

## Acknowledgments

We thank Yi Cui, Francesca Santoro, and Feng Wang for discussions, Osip Schwartz for his help with the initial versions of the multilayer model and the optical set-up, and Edwin Alfonzo for synthesizing electrochromic materials. This work was supported by the David and Lucile Packard Foundation (Cui and Mueller), National Institute of Health (1R01GM125737), The Shurl and Kay Curci Foundation (Kantarci and Zuchero), Stanford Berry Fellowship (Kantarci), the National Science Foundation graduate research fellowships program (NSF-GRFP), Diversifying academia recruiting excellence fellowship program (DARE), and enhancing diversity in graduate education doctoral fellowship program (EDGE) to Alfonso. Part of this work was performed at the Stanford Nano Shared Facilities (SNSF)/Stanford Nanofabrication Facility (SNF), supported by the National Science Foundation under award ECCS-1542152.

## References

1. M. E. J. Obien, K. Deligkaris, T. Bullmann, D. J. Bakkum, U. Frey, Revealing neuronal function through microelectrode array recordings. Front. Neurosci. 8, 423 (2014).

2. B. Miccoli, et al., High-Density Electrical Recording and Impedance Imaging With a Multi-Modal CMOS Multi-Electrode Array Chip. Front. Neurosci. 13, 641 (2019).

3. T.-W. Chen, et al., Ultrasensitive fluorescent proteins for imaging neuronal activity. Nature 499, 295–300 (2013).

4. T. Nagai, K. Horikawa, K. Saito, T. Matsuda, Genetically encoded Ca2+ indicators; expanded affinity range, color hue and compatibility with optogenetics. Front. Mol. Neurosci. 7, 7260 (2014).

5. J. M. Kralj, A. D. Douglass, D. R. Hochbaum, D. Maclaurin, A. E. Cohen, Optical recording of action potentials in mammalian neurons using a microbial rhodopsin. Nat. Methods 9, 90–95 (2011).

6. M. Z. Lin, M. J. Schnitzer, Genetically encoded indicators of neuronal activity. Nat. Neurosci. 19, 1142–1153 (2016).

7. E. W. Miller, Small molecule fluorescent voltage indicators for studying membrane potential. Curr. Opin. Chem. Biol. 33, 74–80 (2016).

8. D. S. Peterka, H. Takahashi, R. Yuste, Imaging voltage in neurons. Neuron 69, 9–21 (2011).

9. W. Akemann, A. Lundby, H. Mutoh, T. Knöpfel, Effect of voltage sensitive fluorescent proteins on neuronal excitability. Biophys. J. 96, 3959–3976 (2009).

10. H. H. Yang, F. St-Pierre, Genetically Encoded Voltage Indicators: Opportunities and Challenges. J. Neurosci. 36, 9977–9989 (2016).

11. D. R. Hochbaum, et al., All-optical electrophysiology in mammalian neurons using engineered microbial rhodopsins. Nat. Methods 11, 825–833 (2014).

12. F. St-Pierre, et al., High-fidelity optical reporting of neuronal electrical activity with an ultrafast fluorescent voltage sensor. Nat. Neurosci. 17, 884–889 (2014).

13. T. H. Grandy, S. A. Greenfield, I. M. Devonshire, An evaluation of in vivo voltage-sensitive dyes: pharmacological side effects and signal-to-noise ratios after effective removal of brain-pulsation artifacts. J. Neurophysiol. 108, 2931–2945 (2012).

14. X.-W. Liu, et al., Plasmonic-Based Electrochemical Impedance Imaging of Electrical Activities in Single Cells. Angew. Chem. Int. Ed Engl. 56, 8855–8859 (2017).

15. Y. Yang, et al., Imaging Action Potential in Single Mammalian Neurons by Tracking the Accompanying Sub-Nanometer Mechanical Motion. ACS Nano 12, 4186–4193 (2018).

16. B. W. Graf, T. S. Ralston, H.-J. Ko, S. A. Boppart, Detecting intrinsic scattering changes correlated to neuron action potentials using optical coherence imaging. Opt. Express 17, 13447–13457 (2009).

17. E. Kreysing, H. Hassani, N. Hampe, A. Offenhäusser, Nanometer-Resolved Mapping of Cell–Substrate Distances of Contracting Cardiomyocytes Using Surface Plasmon Resonance Microscopy. ACS Nano 12, 8934–8942 (2018).

18. G. H. Kim, P. Kosterin, A. L. Obaid, B. M. Salzberg, A mechanical spike accompanies the action potential in Mammalian nerve terminals. Biophys. J. 92, 3122–3129 (2007).

19. T. Ling, et al., Full-field interferometric imaging of propagating action potentials. Light Sci Appl 7, 107 (2018).

20. M. Svrcek, S. Rutherford, A. Y. H. Chen, I. Provaznik, B. Smaill, Characteristics of motion artifacts in cardiac optical mapping studies. Conf. Proc. IEEE Eng. Med. Biol. Soc. 2009, 3240–3243 (2009).

21. J. F. Barry, et al., Optical magnetic detection of single-neuron action potentials using quantum defects in diamond. Proc. Natl. Acad. Sci. U. S. A. 113, 14133–14138 (2016).

22. K. Itaya, K. Shibayama, H. Akahoshi, S. Toshima, Prussian‐blue‐modified electrodes: An application for a stable electrochromic display device. Journal of Applied Physics 53, 804–805 (1982).

23. K. Yamanaka, Anodically electrodeposited iridium oxide films (AEIROF) from alkaline solutions for electrochromic display devices. Jpn. J. Appl. Phys. 28, 632 (1989).

24. J. Hwang, D. B. Tanner, I. Schwendeman, J. R. Reynolds, Optical properties of nondegenerate ground-state polymers: Three dioxythiophene-based conjugated polymers. Phys. Rev. B Condens. Matter 67, 115205 (2003).

25. C. G. Granqvist, Electrochromics for smart windows: Oxide-based thin films and devices. Thin Solid Films 564, 1–38 (2014).

26. Z. D. Marks, et al., Switchable diffractive optics using patterned PEDOT:PSS based electrochromic thin-films. Organic Electronics 37, 271–279 (2016).

27. M. F. Zainal, Y. Mohd, Characterization of PEDOT Films for Electrochromic Applications. Polymer-Plastics Technology and Engineering 54, 276–281 (2015).

28. Y. Wen, J. Xu, Scientific Importance of Water-Processable PEDOT--PSS and Preparation, Challenge and New Application in Sensors of Its Film Electrode: A Review. J. Polym. Sci. A Polym. Chem. 55, 1121–1150 (2017).

29. Z. Aqrawe, J. Montgomery, J. Travas-Sejdic, D. Svirskis, Conducting polymers for neuronal microelectrode array recording and stimulation. Sens. Actuators B Chem. 257, 753–765 (2018).

30. S. J. Wilks, S. M. Richardson-Burns, J. L. Hendricks, D. C. Martin, K. J. Otto, Poly(3,4-ethylenedioxythiophene) as a Micro-Neural Interface Material for Electrostimulation. Front. Neuroeng. 2, 7 (2009).

31. G. Dijk, A. L. Rutz, G. G. Malliaras, Stability of PEDOT:PSS‐Coated Gold Electrodes in Cell Culture Conditions. Adv. Mater. Technol. 5, 1900662 (2020).

32. C. Boehler, Z. Aqrawe, M. Asplund, Applications of PEDOT in bioelectronic medicine. Bioelectronics in Medicine 2, 89–99 (2019).

33. P. Simon, Y. Gogotsi, B. Dunn, Where do batteries end and supercapacitors begin? Science (2014).

34. A. A. Argun, et al., Multicolored Electrochromism in Polymers: Structures and Devices. Chemistry of Materials 16, 4401–4412 (2004).

35. J. Kawahara, P. A. Ersman, I. Engquist, M. Berggren, Improving the color switch contrast in PEDOT:PSS-based electrochromic displays. Organic Electronics 13, 469–474 (2012).

36. D. Levasseur, I. Mjejri, T. Rolland, A. Rougier, Color Tuning by Oxide Addition in PEDOT:PSS-Based Electrochromic Devices. Polymers 11 (2019).

37. W. Shi, et al., Micron-thick highly conductive PEDOT films synthesized via self-inhibited polymerization: roles of anions. NPG Asia Materials 9, e405–e405 (2017).

38. M. Ouyang, et al., Enhanced electrochromic switching speed and electrochemical stability of conducting polymer film on an ionic liquid functionalized ITO electrode. New J. Chem. 39, 5329–5335 (2015).

39. S. Hassab, et al., A new standard method to calculate electrochromic switching time. Sol. Energy Mater. Sol. Cells 185, 54–60 (2018).

40. T. Xu, et al., High-contrast and fast electrochromic switching enabled by plasmonics. Nat. Commun. 7, 10479 (2016).

41. D. A. Koutsouras, et al., Impedance Spectroscopy of Spin-Cast and Electrochemically Deposited PEDOT:PSS Films on Microfabricated Electrodes with Various Areas. ChemElectroChem 4, 2321–2327 (2017).

42. E. Stavrinidou, et al., Direct measurement of ion mobility in a conducting polymer. Adv. Mater. 25, 4488–4493 (2013).

43. H.-E. Tseng, T.-H. Jen, K.-Y. Peng, S.-A. Chen, Measurements of charge mobility and diffusion coefficient of conjugated electroluminescent polymers by time-of-flight method. Applied Physics Letters 84, 1456–1458 (2004).

44. V. V. Fedorov, et al., Application of blebbistatin as an excitation–contraction uncoupler for electrophysiologic study of rat and rabbit hearts. Heart Rhythm 4, 619–626 (2007).

45. C. Gold, D. A. Henze, C. Koch, G. Buzsáki, On the origin of the extracellular action potential waveform: A modeling study. J. Neurophysiol. 95, 3113–3128 (2006).

46. M. L. Fiszman, J. L. Barker, S. V. Jones, Electrophysiological responses to muscarinic receptor stimulation in cultured hippocampal neurons. Brain Res. 557, 1–4 (1991).

47. S. Haj-Dahmane, R. Andrade, Muscarinic activation of a voltage-dependent cation nonselective current in rat association cortex. Journal of Neuroscience (1996).

48. H. Cho, J. Shin, C. Y. Shin, S.-Y. Lee, U. Oh, Mechanosensitive ion channels in cultured sensory neurons of neonatal rats. J. Neurosci. 22, 1238–1247 (2002).

49. J. N. Wood, et al., Capsaicin-induced ion fluxes in dorsal root ganglion cells in culture. J. Neurosci. 8, 3208–3220 (1988).

50. W. Greffrath, S. T. Schwarz, D. Büsselberg, R.-D. Treede, Heat-induced action potential discharges in nociceptive primary sensory neurons of rats. J. Neurophysiol. 102, 424–436 (2009).

51. J. B. Zuchero, Purification of dorsal root ganglion neurons from rat by immunopanning. Cold Spring Harb. Protoc. 2014, 826–838 (2014).

52. W. Zhu, G. S. Oxford, Differential gene expression of neonatal and adult DRG neurons correlates with the differential sensitization of TRPV1 responses to nerve growth factor. Neurosci. Lett. 500, 192–196 (2011).

53. K. Newberry, et al., Development of a spontaneously active dorsal root ganglia assay using multiwell multielectrode arrays. J. Neurophysiol. 115, 3217–3228 (2016).

54. N. Farah, et al., Holographically patterned activation using photo-absorber induced neural–thermal stimulation. J. Neural Eng. 10, 056004 (2013).

55. S. Yoo, J.-H. Park, Y. Nam, Single-Cell Photothermal Neuromodulation for Functional Mapping of Neural Networks. ACS Nano 13, 544–551 (2019).

56. N. Martino, et al., Photothermal cellular stimulation in functional bio-polymer interfaces. Sci. Rep. 5, 8911 (2015).

57. A. Greppmair, et al., Measurement of the in-plane thermal conductivity by steady-state infrared thermography. Rev. Sci. Instrum. 88, 044903 (2017).

58. S. Wäldchen, J. Lehmann, T. Klein, S. van de Linde, M. Sauer, Light-induced cell damage in live-cell super-resolution microscopy. Sci. Rep. 5, 15348 (2015).

59. C. Fourie, M. Kiraly, D. V. Madison, J. M. Montgomery, Paired whole cell recordings in organotypic hippocampal slices. J. Vis. Exp., 51958 (2014).

60. J. G. Rohan, K. A. Carhuatanta, S. M. McInturf, M. K. Miklasevich, R. Jankord, Modulating Hippocampal Plasticity with In Vivo Brain Stimulation. Journal of Neuroscience 35, 12824–12832 (2015).

61. J. H. Williams, J. A. Kauer, Properties of carbachol-induced oscillatory activity in rat hippocampus. J. Neurophysiol. 78, 2631–2640 (1997).

