## Supplementary material for "Label-free optical detection of bioelectric potentials using electrochromic thin films": SI

### Supplementary Information

### Supplementary Text

**Section 1: Modeling the light reflection at the glass:ITO:PEDOT:water interface**

A theoretical study of the reflectivity of the PEDOT:PSS layer can help us to understand how to maximize the signal-to-noise ratio of optically detecting a given voltage change $\Delta V$. The basic physics is reflection at the boundary between media having different refractive indices $\underline{n}=n+i\kappa$, where $n$ is the usual refractive index and $\kappa$ the extinction coefficient (1).

Applied voltages change the absorptivity or extinction coefficient of the PEDOT:PSS layer by $\Delta\kappa = (d\kappa/dV)\Delta V$, which in turn change the reflectivity of the layer by $\Delta R = (dR/d\kappa)\Delta\kappa$. Background signals from technical sources (such as vibrations, air currents, or dust particles) are generally proportional to the detected power, and thus to the reflectivity $R$. To maximize the signal-to-noise-ratio, we maximize $\Delta R/R = S\Delta\kappa$, where $S = (1/R)(dR/d\kappa)$. If $S \gg 1$, the reflectivity change is optically enhanced relative to the change in $\kappa$.

Our model is a four-layer structure (Fig. 1C) in which light enters from glass BK-7 ($n_0 = 1.51$ at 660 nm and incident angle $\theta_0$) into the ITO layer ($n_1 = 1.84 + i0.061$, thickness $d_1$, incident angle $\theta_1$), then into the PEDOT:PSS layer ($n_2 = 1.4 + i\kappa$, thickness $d_2$, incident angle $\theta_2$), and finally into water ($n_3 = 1.33$, incident angle $\theta_3$). The total reflectivity of the multilayer system is determined by interference between reflections from all layers. It can be calculated using the transfer matrix method. We define the wavenumber $k = \omega/c$ in vacuum as well as the *z*-components of the wavenumbers in material, $k_{i,z} = kn_i \cos(\theta_i)$, where $n_i$ and $\theta_i$ are the index of refraction and incident angle in the $i^{th}$ layer. The angles can be calculated iteratively from $\theta_0$ by applying $\theta_{i+1} = \arcsin[n_i \sin(\theta_i)/n_{i+1}]$. They must be allowed to be complex, given that some of the $n_i$ are complex. The (amplitude) reflection coefficients for a beam in medium *i* going into medium *j* are given by

$r_{ij}^p = (n_j \cos \theta_j - n_i \cos \theta_i)/(n_j \cos\theta_j + n_i \cos \theta_i)$ (p-polarization),

$r_{ij}^s = (n_j \cos \theta_i - n_i \cos \theta_j)/(n_j \cos\theta_i + n_i \cos \theta_j)$ (s-polarization).

In the experiment, we use s-polarization. We see that it leads to low background reflectivities and thus a high contrast $\Delta R/R$ for a given electrical potential difference. For each boundary between layer *i* and *i*+1 (i=0, 1, and 2) we define a matrix

$$M_i = \begin{pmatrix} \exp(-i\delta_i) & r_{ij}\exp(-i\delta_i) \\ r_{ij}\exp(i\delta_i) & \exp(i\delta_i) \end{pmatrix},$$

where $\delta_0 = 0$ and $\delta_i = d_i k_{i,z}$ for $i$=1 and 2. To obtain the reflectivity of the entire system, we calculate the product

$$M = \frac{1}{t_{01}} M_0 M_1 M_2 \equiv \begin{pmatrix} M_{11} & M_{12} \\ M_{21} & M_{22} \end{pmatrix},$$

where $t_{01}$ is the transmission coefficient at the first boundary. The reflectivity of the entire system is the ratio of the elements in the first column of the matrix, $r = M_{21}/M_{11}$. The absolute square $R = |r|^2$ gives the power reflectivity.

The angle $\theta_0$ is the incident angle of the beam from the prism into ITO. It is related to the incident angle $\theta$ of the beam going into the prism by $\theta_0 = \arccos[\cos(\beta/2 + \theta)/n_0] - \beta/2$, where $\beta$ is the apex angle of the prism, 60° in our case.

For vanishing ITO and PEDOT:PSS thicknesses, $d_1 = d_2 = 0$, the model reproduces the results of simple theory, such as the onset of total internal reflection at an angle of $\theta_0 = \arcsin[n_3/n_0] = 61.7°$ (which corresponds to $\theta = 62.3°$ measured outside the prism).

We compare the model to observation at a wavelength of $\lambda = 660$ nm. To validate the model, we compare the measured and predicted reflectivities as a function of angle $\theta$ for various PEDOT:PSS thicknesses (Fig. S3), obtaining good qualitative agreement. The shape of the curves obtained from the model, i.e. the reflectivity, first increasing as the angle increases and then decreasing sharply after the total internal reflection angle, closely follows the experimental measurements. The samples were prepared with thicknesses of 30, 60, and 100 nm, but the PEDOT:PSS layer thickness can be inhomogeneous, and so the exact thickness at the point probed by the beam has a relatively high variance. We thus determine the thickness of the PEDOT:PSS layer to be 34, 59, and 77 nm by fitting the measured reflectivity (Fig. S3).

We also measured the dependence of the reflectivity on the applied bias potential. Figure S4A shows a measurement of the reflectivity of a PEDOT:PSS film as a function of the bias voltage of the data. Matching the theoretically calculated reflectivity (with $d_1$ =30 nm ITO and $d_2$ =120 nm PEDOT:PSS) with the measured one allows us to calculate the extinction coefficient $\kappa$ as a function of voltage (Fig. S4B). Mathematically, for each measured reflectivity, this yields two solutions for $\kappa$, of which only one is physical. When using PEDOT:PSS to sense action potentials, we do not apply any external bias and the layer adopts an open-circuit potential of -50 mV. At this operating point, we find $\kappa_0 \sim 0.22$ and $d\kappa/dV = -0.57$/V.

We also use the model to predict the signal strength $|\Delta R/R|$ of the optical readout as a function of the bias voltage on the PEDOT:PSS layer (Fig. S4C) as well as the incident angle (Fig. S4D). For a PEDOT:PSS layer that is $d_2$ = 120 nm thick, for a 1 mV change, we predict that the sensitivity $|\Delta R/R|$ will have a global maximum at a certain bias voltage and then decreases as the bias voltage is scanned in either direction. At about 100 mV, $\Delta R/R$ has a zero crossing; at higher bias potential, $|\Delta R/R|$ will increase again. This prediction is verified in detail by the experimental data (Fig. S4C). The modeled signal strength as a function of angle is compared with the data shown in Fig. 2a (Fig. S4D). It peaks at an incident angle $\theta = 67^o$ (measured outside the prism), showing good agreement between the model and the data.

Note that the measured fractional reflectivity changes $\Delta R/R$ can be larger than the changes in the extinction coefficients $\Delta\kappa$ by as much as ten-fold, i.e., $S = (1/R)(dR/d\kappa) \gg 1$. Such optical enhancement is a key factor in the sensitivity of ECORE. Our model predicts that S can reach values of 100, indicating that there is much room for optimizing the sensitivity of ECORE further.

### Section 2: The electrochromism of PEDOT

3,4-ethylenedioxythiophene (EDOT) is a colorless, viscous compound which upon oxidative polymerization forms the better known insoluble blue solid poly-(3,4-ethylenedioxythiophene) (PEDOT). This organic polymer belongs to the class of conductive polymers due to its ability to support electronic and ionic transport, which are not typically displayed by their insulating organic counterparts. Electronic conductivity is achieved by the extended π-conjugated system along the backbone of the polymer. Extended conjugated electron systems like dyes and pigments typically absorb in the visible light range of the optical spectrum corresponding to a π-π* and/or a nonbonding-π* electronic transition; the same extended conjugated system gives rise to PEDOT's color.

Upon oxidation of the polymer backbone, an electron is removed from the π-conjugated system. This newly created positive charge is stabilized by the delocalization of the charge over several EDOT constituents and the recruitment of anions from the electrolyte solution. Doping or introduction of poly(styrene sulfonic acid) (PSS) impurities during the synthesis improves the water processability of the polymer and assists in the stabilization of the positive charge by providing an excess of negative charge from the sulfonate groups but requiring cations from the electrolyte for charge compensation. The effect of twisting present in all polymers in solution restricts the length of the conjugated system creating a distinct localized charge state known as a polaron. The creation of a polaron alters the optical properties of the polymer by creating a lower energy state available in which optical transitions can occur at a lower energy cost thus leading to a greater absorption of lower energy light in the near-infrared range of the electromagnetic spectrum. The electrochromism of PEDOT:PSS originates from the low charge or potential required to reversibly remove an electron from the backbone of the polymer thus affecting the optical transitions available due to the creation of the polaron.

Images of the electrodeposited film obtained through scanning electron microscopy display a flat surface with granular domains on the order of 100 nm (Supplementary Fig. 1). Visual inspection of the electrodeposited PEDOT:PSS film shows that the film appears blue at positive potentials such as +500 mV, while it appears purple at negative potentials such as -500 mV (Supplementary Fig. 2). The optical properties of electrodeposited PEDOT:PSS films are different from those of spin-coated PEDOT:PSS films from pre-formed polymer solutions. For instance, in the reduced state, electrodeposited PEDOT:PSS has a maximum absorbance at $\lambda_{max}$ = 580 nm while the spin-coated film has a maximum absorbance at $\lambda_{max}$ = 650 nm (2). This red-shifted optical spectrum of the spin-coated film can be attributed to a longer conjugation length and increased planarity of the pre-formed PEDOT:PSS polymer chains through chemically oxidative polymerization (3).

### Section 3: Spatial-temporal response of PEDOT

#### Temporal Resolution

The temporal response of the film must also be sufficiently fast in order to capture cellular electric potentials in the order of 1 ms. The limit on the temporal resolution is given by the $\tau$ = RC product of the electrolyte resistance $R$ and the double-layer capacitance $C$ of the PEDOT:PSS film. Electrochemical impedance spectroscopy (Fig. S5) measures the resistance and capacitance properties of an electroactive material via application of a sinusoidal AC voltage signal. A 10-mV sinusoidal signal with a DC bias of 0 mV vs. Ag/AgCl was used to measure the impedance $Z$ of films deposited on patterned ITO in the frequency range of 100 mHz to 100 kHz. Figure S5 shows the impedance $|Z|$ as a function of the frequency for PEDOT samples of different surface areas. The results were fitted with a resistor-capacitor series connection $Z = R+1/(i\omega C)$, where $\omega$ is the angular frequency, using the EC-Lab software (Bio-Logic), where $R$ represents the electrolyte resistance and $C$ the double layer capacitance. At low frequencies, Z is dominated by the capacitive term $1/(i\omega C)$ and thus proportional while at high frequencies, the constant resistance $R$ dominates. Thus, the data allows us to independently determine $R$ and $C$ (Table S1). The double layer capacitance $C = (8.11\pm0.44)$ µF/mm$^2 \times A$ is found to be proportional to the PEDOT surface area $A$ as expected. The resistance is expected to be proportional to $1/A^{1/2}$ if the PEDOT surface area is small compared to the dimensions of the rest of the system (4) . Our measurements reproduce this, with R = (532±15) Ω mm/$A^{1/2}$, in good agreement with expectations from the specific resistance of the medium of 69 Ω cm for a 0.9% saline solution at 22°C. Since $R$ and $C$ are proportional to $1/A^{1/2}$ and $A$, respectively, the $RC$ time constant is proportional to $A^{1/2}$, i.e., it decreases with the square-root of the surface area when the surface area is decreased. The 10%-90% rise-time for square-wave signals is $t_r$ =2.2$RC$; we find it to be $t_r$ =9.5 ms mm/$A^{1/2}$ (Table. S1).

**Spatial resolution**

The spatial resolution of the PEDOT:PSS film must be sufficient to detect electrical activity from individual cells. When a cell is in close contact with the film, the cell's electrical activity will charge the film locally; however, the PEDOT:PSS film is electrically conductive and the charge diffusion may degrade the spatial resolution. The reported charge mobility in PEDOT polymer is around $10^{-4}$ to $10^{-6}$ $cm^2/V/s$ (5, 6), which implies a diffusion constant $D = k_B T\mu/e$ between $2.5\times10^{-3}$ and $2.5\times10^{-1}$ $\mu m^2/ms$. After 1 ms, the diffusion area will be between $10^{-3}$ and $10^{-1}$ $\mu m^2$, much smaller than a cell. Nevertheless, we experimentally measured the spatial resolution of the film by measuring the optical response at different film locations when a 20 mV electrical pulse of 50 ms duration is applied through a microelectrode 10 μm above the film. The measured full width at half maximum for the optical response is 33.4 μm, which is the size of the laser spot (Fig. 2f). The optical response decayed quickly as the microelectrode was moved away from the laser spot. Therefore, the spatial resolution of the electrochromic film is limited not by charge diffusion, but by the size of the probing laser spot.

### Materials and Methods

#### Electrodeposition, cyclic voltammogram, and electrochemical impedance characterization of PEDOT:PSS (referred as PEDOT hereafter) films

A 2% aqueous solution of poly(sodium 4-styrene sulfonate) (Mw = 70,000) (w/w%) was mixed with 10 μL of 3,4 ethylenedioxythiophene (EDOT) to make a 10 mM solution. Prior to deposition, ITO-coated unpolished float glass (Rs = 70-100 Ω/sq, 15-30 nm ITO thickness) from Delta Technologies were cleaned using UV-Ozone. The monomer was electropolymerized by applying a constant voltage of 950 mV on ITO vs. Ag/AgCl resulting in the polymer PEDOT on the ITO surface. The thickness of the PEDOT film was measured using a Bruker Dektak XT profilometer. Cyclic voltammogram measurements of the PEDOT film were taken with a scan rate of 50 mV/s restricted by the potential window of -500 mV to +500 mV vs Ag/AgCl in HEPES buffered Tyrode's salt solution.

#### ECORE Optical instrumentation

The ECORE optical setup consisted of PEDOT:PSS ($n_{PEDOT:PSS}$ =1.4 @ 660 nm) thin layer deposited onto ITO ($n_{ITO}$ =1.7 @ 660 nm) glass. The sample was mounted onto a translational stage holding a BK-7 ($n_{BK7}$ =1.5142 @ 660 nm) equilateral prism. A thin layer of Type F immersion oil ($n_{oil}$ =1.5124 @ 660 nm) was applied to the bottom of the ITO glass for index matching with the prism and suppression of the back reflection from the ITO-coated glass. A 20-mW, 660-nm laser diode (ThorLabs LP633-SF50), pigtailed with a single-mode fiber, was mounted on a temperature controlled laser diode mount (Thorlabs, Inc.) equipped with a current and temperature controller. The laser beam polarization was cleaned up by a combination of two half-wave plates, a broadband polarizing beamsplitter cube and a Wollaston prism. The Wollaston prism separated polarized light into two orthogonal linearly polarized outgoing beams. The amount p- and s-polarized light was manually controlled by the preceding half-wave plate. The balancing of the light intensity consisted of manually rotating the half-wave plate.

The beam spot size was determined by fitting the intensity profile of the laser beam captured by a CCD camera with a Gaussian function. The measured full width at half maximum for the beam spot was 33.4

μm. The average intensity (W/cm$^2$) over the aperture that transmits ~99% of the power was obtained as $P_0/(2\pi\,(FWHM)^2)$ from a measurement of the power ($P_0$) using a silicon based power meter.

The sample and the reference beam were redirected to the two diodes of a homemade differential photodiode detector. The output was amplified using an amplifier (Axon Instruments Axopatch-1D Patch Clamp amplifier) and digitized at 10 kHz using Axon Digidata 1440A low noise digitizer.

**Patterning of the ITO through lithography to control the area of PEDOT film.**

ITO glass was cleaned using Samco ozone cleaner which used a UV producing light bulb and a molecular ozone source to generate ozone and remove organic molecules on the substrate surface. Following the cleaning, the surface was coated with hexamethyldisilazane (HMDS) using the YES prime oven. Shipley 3612 positive resist was spin coated at 5.5K RPM for 30 seconds to produce a 1 μm thick film. The photoresist is prebake at 90$^o$C for 1 minute. The pattern was designed and transferred onto the substrate using Heidelberg maskless aligner (MLA150). After exposure to square patterns with areas of either 9 mm$^2$,4 mm$^2$,1 mm$^2$,0.56 mm$^2$, and 0.25 mm$^2$, the film was post exposure baked at 115$^0$C for 1 minute. Then, the substrate was developed using MEGAPOSIT$^{TM}$ MF$^{TM}$ -26A developer and rinsed with water. The patterned ITO substrates were used to electrodeposited PEDOT:PSS films with defined film areas.

**Optimization of incident angle, thickness, bias potential**

Commercial ITO substrates were patterned using standard photolithography techniques to define the surface area of interest. PEDOT films were electrodeposited onto the patterned ITO substrate. Using a potentiostat (Bio-Logic, SP200) and a 3-electrode cell setup, the film was subjected to periodic electric pulses with an amplitude of $\Delta V$ = 1 mV, at a constant bias $V_0$ = 0 mV, and a frequency of 10 Hz, and the measured $\Delta R/R$ peak-to-peak amplitude recorded. This method was used during the optimization of the incident angle and film thickness. The prism attached to a rotation stage was adjusted manually during the optimization of the incident angle. Different film thickness was acquired by holding the potential (950 mV) constant and limiting the charge consumed during the electrodeposition. For the optimization of the bias potential $V_0$ the film is subjected to periodic electric pulses with an amplitude of $\Delta V$ = 1 mV, and a frequency of 10 Hz. Finally, at $V_0$=0, the optical response of the PEDOT system is characterized by applying square-waves of varying amplitudes. Due to hardware limitations of the potentiostat, we are unable to apply electric pulses smaller than 1 mV.

**Preparation of cultured monolayer of cells: stem cells derived cardiomyocytes, and Primary hippocampal neurons**

Cryopreserved hiPSC-CM (Cor.4U [Axiogenesis AG, Cologne, Germany] were kept in liquid nitrogen until culture.  The cells were thawed and plated according to the instructions provided by the manufacturer. The cells were cultured in Geltrex$^{TM}$ treated PEDOT:PSS film in a humidified incubator at 37$^o$C. The subsequent maintenance protocols followed manufacturer's instructions and use of corresponding maintenance media (Cor. 4U-maintenance media) supplemented with ciprofloxacin (2 mg/ mL). Experiments were performed between days 4-7 after plating as recommended by the manufacturer. All plated cells showed regular contractions after 3h of the initial plating. During optical measurements of the electric potential, the maintenance medium was replaced with modified Tyrode solution (4.2 mM $K^+$ instead of the standard 2.7 mM $K^+$).

Primary embryonic rat Hippocampal neurons were isolated from Sprague-Dawley fetal rats (age E-15-E16) (7). Briefly, hippocampal neurons of rat embryos were dissected in Hank's buffer solution and enzymatically treated in 0.25% trypsin at 37$^o$C for 30 minutes followed by mechanical dissociation by passing through a fire polished Pasteur pipette for less than 10 minutes. Dulbecco's modified Eagle medium (DMEM) containing 15% FBS was used to stop the trypsinization process and cells were spun down at the bottom of the collection tube. Dissociated Hippocampal neurons were resuspended, counted and plated on either culture dish or the PEDOT:PSS film coated with poly-D-lysine. Cultures were maintained in a neurobasal medium supplemented with B27. All cultures were housed in a humidified incubator at 37$^o$C supplied with 5% $CO_2$ and neurons were selected by applying 4 μM cytosine arabinoside (1-b-D-arabinofuranosylcytosine) for 24 hours to cultures 2 days after plating cells. This method produces neuronal cultures that are free of non-neuronal cells.

**Purification of primary embryonic rat dorsal root ganglion neurons (DRGs).**

DRGs were performed by a method adapted from that described in Zuchero's publications (8, 9).

Preparation of samples for plating neurons:

1 mg/ml of Poly-D-Lysine hydrobromide stock (Sigma-Aldrich, Cat#P6407) was diluted 100-fold in sterile ddH2O and added to plates for 30 mins at room temperature. Plates were rinsed with sterile ddH2O three times and coated with 1 mg/ml Laminin (R&D Systems Cat#3400-010-02) diluted 1:200 in neurobasal media (Thermo Fisher Scientific, Cat# 21103049). Laminin coating was done at 37 °C for 4 to 24 hours. Laminin was removed from the plates immediately before plating neurons and replaced with DRG base media to prevent drying.

Preparation of immunopanning plates:

BSL-1 plate: One 15 cm petri dish was coated with 40 ml of 2 mg/ml BSL-1 (Vector Labs, Cat #L-1100) in 20 ml PBS overnight at 4 °C. The plate was rinsed three times with sterile PBS and coated with 9 ml of 0.2% BSA (Sigma-Aldrich, Cat#A-8806) at room temperature for at least 2 hours before using.

CD9 plate: One 15 cm petri dish was coated with 90 ml of goat-anti-mouse IgG+ IgM (H+L) secondary antibody (Jackson ImmunoResearch, Cat# 115-005-044) in 20 ml of Tris-HCl (pH 9.5) overnight at 4°C.The plate was rinsed three times with sterile PBS and coated with anti-rat CD9 antibody (Thermo Fisher Scientific, #BDB551808) diluted 1:200 in 0.2% BSA.

Dissections: All animal procedures were approved by the Stanford University's Administrative Panel on Laboratory Animal Care. Two timed pregnant Sprague-Dawley rats (Charles Rivers) with E15 embryos were euthanized by $CO_2$ inhalation. The placentas were dissected out and placed in Leibovitz's L15 dissection medium (Thermo Fisher Scientific, 11415114) supplemented with 10% FCS (Thermo Fisher Scientific, A3160401). Embryos were carefully removed from the amniotic sacs and placed in a separate 10 cm dish with the dissection medium.

Following decapitation, all of the organs and the spinal column on the ventral side of the embryos were carefully removed until the spinal cord was entirely exposed along the anteroposterior axis. This step dissociates the spinal cord from remaining ventral tissues and allows separation and visualization of DRGs. The embryos were then flipped so that the dorsal side was facing up. The skin above the spinal cord was peeled off with two forceps revealing DRGs lateral to the spinal column across its entire length. DRGs were then cut out at their bases using small scissors. 20-30 embryos were pooled for each prep, and each prep was treated as one biological replica.

Immunopanning:

Dissected DRGs were manually collected into D-PBS. The immunopanning protocol was performed essentially as described in Zuchero's publication (8, 9). Briefly, dissociated cells were passed serially over a BSL-1 plate (to remove blood and endothelial cells) and a CD9 plate (to remove glial cells). This protocol yields a single-cell suspension of 1.5-2 million DRGs with higher than 98% purity, devoid of contaminating glia, blood cells or fibroblasts.

Plating and Culturing:

Isolated DRG neurons were resuspended at 1000 cells/ml volume and plated in densities ranging from 10,000 to 60,000 cells per plate. Neurons were placed on the bottom of the plates containing 1 ml of DRG base media to achieve dispersed plating. DRG base medium (Zuchero, 2014a, 2014b) was supplemented with 100 ng/ml NGF (Neuromics, Cat#gt15057), 50 ng/ml BDNF (PeproTech, Cat#450-02) and 1 ng/ml NT3 (PeproTech, Cat#450-03) before each feeding (half volume media change) every 2-3 days.

**Preparation of hippocampal brain slices**

All animal procedures conformed to the NIH guide for the Care and Use of Laboratory Animals and were approved by the Stanford University Administrative Panel on Laboratory Animal Care. Hippocampal slices from Sprague-Dawley rat pups were prepared according to the protocol established by C. Fourie *et al.* (*33*). Briefly, the brain slice culture dishes were prepared before the animal dissection by placing 1 mL of culture media into each 35 mm culture dish and carefully putting the membrane insert at a roughly 45 degree angle to the culture media to avoid air bubbles being added under the membrane insert. Six to eight of the 35 mm culture dishes were placed into one 150 mm petri dish followed by a transfer to a $CO_2$ incubator for a 1 hour incubation prior to the dissection. Rat Pups at Postnatal Day 7 (P7) were euthanized by rapid decapitation. Afterwards, the brain was removed and placed into a chilled dissection medium. The cortex was removed from the midbrain to expose the hippocampus. The isolated hippocampus was removed and trimmed from the rest of the brain. Transfer the hippocampi into a new dish containing chilled dissection medium. Using a manual tissue slice to slice the hippocampi transversely into 400 μm sections. The hippocampal slices were transferred into dissection medium and inspected under a dissection microscope for the purpose of removing damaged brain slices. Place 3 to 4 individual slices onto a membrane filter insert in the culture dishes that were pre-warmed in the incubator. The slices were maintained in a 5% $CO_2$ incubator at 37$^o$C and exchanged the medium on the second day. Make sure to pre-warm the culture medium before exchange: add 1 mL of fresh culture medium to each culture dish and place the culture dishes in the $CO_2$ incubator for at least one hour. Transfer the membrane insert into a pre-warmed petri dish containing culture medium. Repeat these steps on the third day after making cultures and then transfer the cultures to an incubator set at 34$^o$C. The medium was changed twice each week and maintained *in vitro* for 7-21 days before recording.

**Electrical and Optical measurements of cultured monolayer of cells and hippocampal brain slices**

For the electrical measurements, a low-noise amplifier with sixty channels (MEA1060-Inv-BC, *Multi Channel Systems MCS GmbH*) was used for the electrophysiology measurements with a sampling rate of 5 kHz. A Ag/AgCl pellet electrode (E200, *WarnerInstruments*) grounded the amplifiers in the bath medium.

All measurements with the cultured monolayer cells were made in a tyrode solution initially heated to 37$^o$C. PEDOT-ITO glass was sterilized using a solution of 70% ethanol. Afterwards, the film was coated with fibronectin, Geltrex™, or Poly-D-lysine to promote the adhesion of the cells onto the surface. Optical measurements of the device were taken after 3-5 days after cell plating with the exception of neuronal cells which were plated up to 4 weeks.

Optical measurements with brain slices were made in an ACSF solution initially heated to 37ºC. After sterilization, the film was coated with Poly-D-lysine to promote the adhesion of the cells onto the surface. The PEDOT film was incubated in the cell incubator overnight with cell medium to allow for sufficient time for the reduction of the PEDOT by the cell medium from blue to purple state. The hippocampal brain slices were cut from the millicell insert and placed upside down onto the surface and gentle force was applied using a 3D-printed adapter terminated with a nylon mesh.

### Supplementary Figures

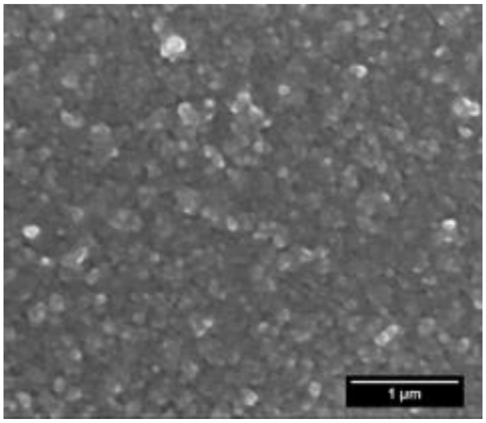


**Fig. S1.**

Scanning electron microscope image of the electrodeposited PEDOT:PSS film.

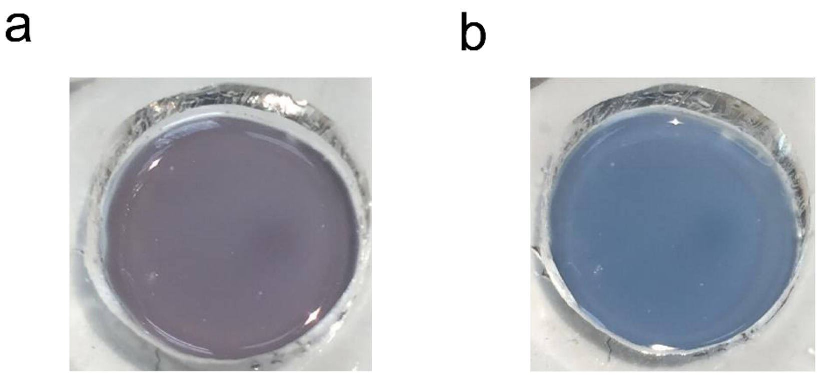


**Fig. S2.**
Photographic image of the electrodeposited PEDOT:PSS film in the a) reduced state and b) oxidized state.

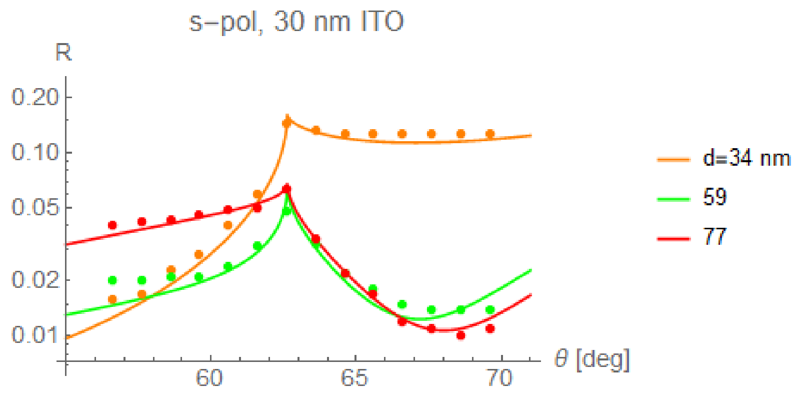


**Fig. S3.**

Measured reflectivity (dots) compared with theory (lines) as a function of the incident angle θ of light into the prism for four different thicknesses of the PEDOT:PSS film. Note that the PEDOT:PSS sample used here was in the pristine state with a higher extinction coefficient κ ~0.4, while voltage sensing is usually performed with zero bias or -50 mV open circuit where PEDOT having κ~0.2.

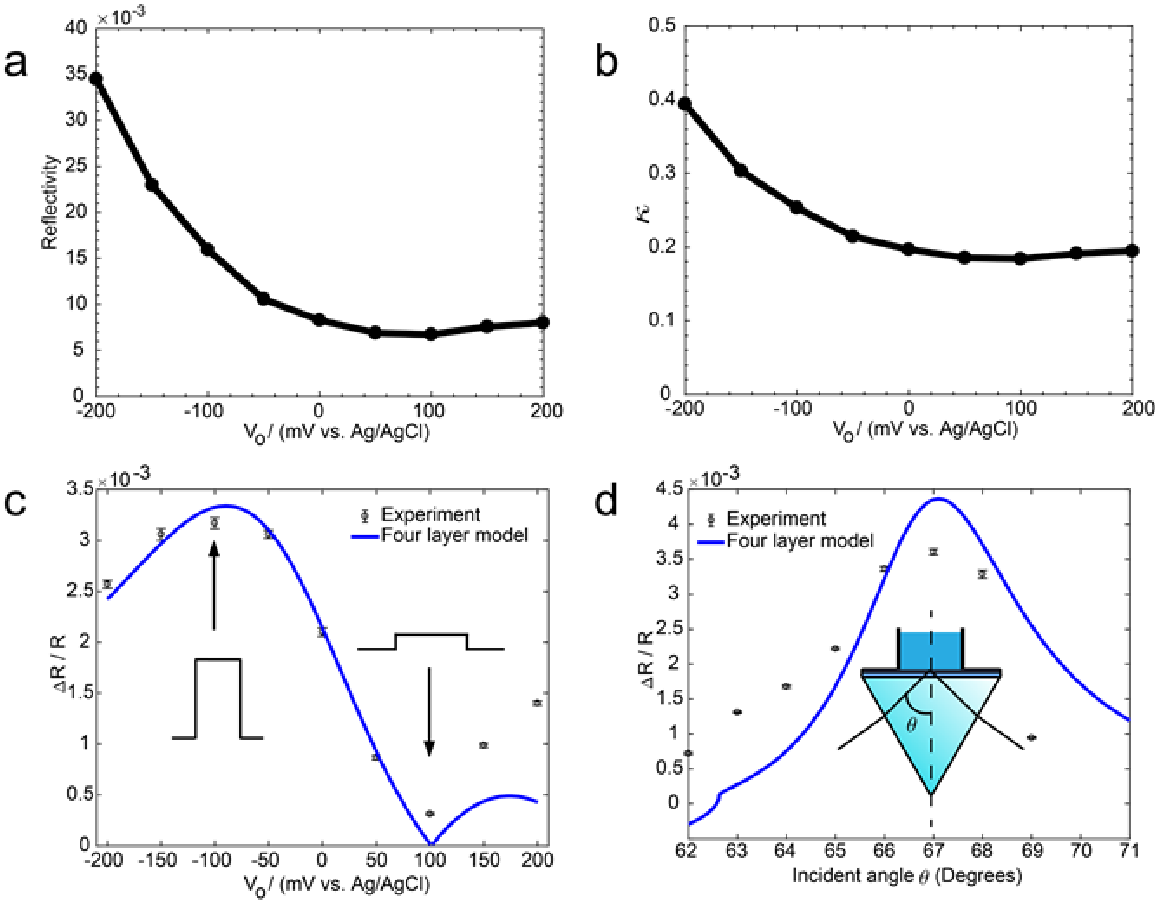


**Fig. S4.**

A) Measured reflectivity as a function of bias, for the same sample used for Fig. 2c. B) Extinction coefficient κ as a function of bias voltage inferred from this data. Dots have been derived from the measurements shown in (A) while the solid line represents a fit, $\kappa = 0.204 - 0.291V + 2.433V^2 - 4.890V^3$, where $V$ is the applied potential in Volt. C) Fractional reflectivity change $|\Delta R/R|$ for a 1-mV applied square waves signal as a function of bias electrode potential. The dots are the measurement from Fig. 2c in the main manuscript. The blue line is the prediction of our model, using the extinction coefficient as determined in (B). D) Fractional reflectivity change $|\Delta R/R|$ for a 1-mV applied square waves signal as a function of incident angle (from Fig. 2a) compared to theory (blue line).

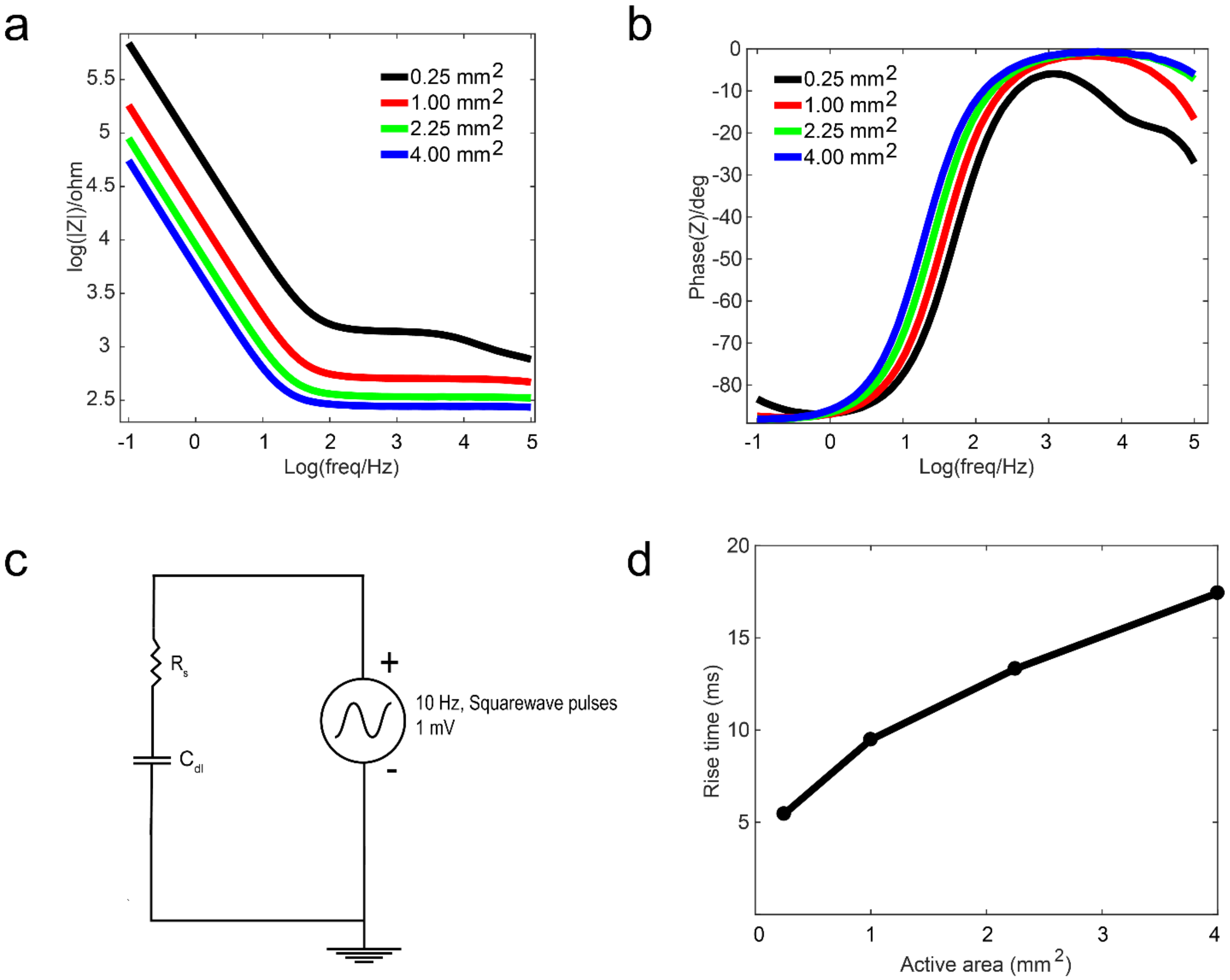


**Fig. S5.**

Impedance spectroscopy of the PEDOT thin films, a) Bode plot demonstrating an increase in the impedance with a decrease in the surface area b) phase angle c) equivalent circuit model used to estimate the expected rise time d) expected rise time as a function of active area.

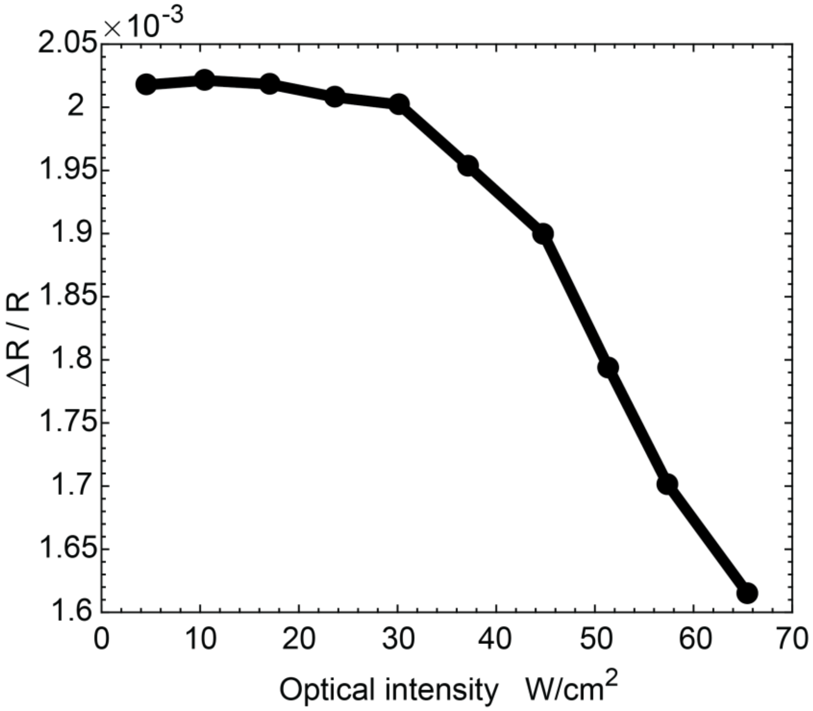


**Fig. S6.**

Fractional reflectivity change $|\Delta R/R|$ for a 1-mV applied square waves signal as a function of optical intensity in W/cm$^2$.

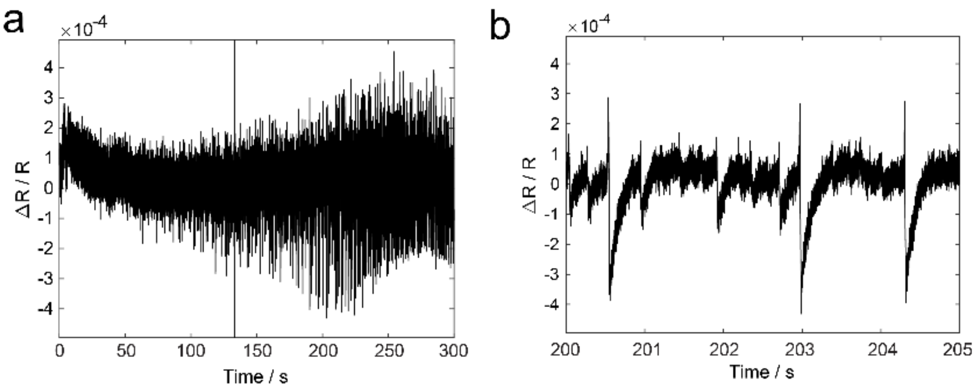


**Fig. S7.**

Hippocampal brain slices exposed to carbachol. a) Recovery of electrical spikes after re-saturation of the ACSF solution by carbogen, b) close-up of the biphasic optical spikes.

### Supplementary Table

#### Table S1.

Temporal dependence on the response time as a function of surface area.

| Surface Area ($mm^2$) | $R_{solution}$ (Ω) | $C_{dl}$ (μF) | Rise time (10%-90%) calculated from RC (ms) | Rise time (10%-90%) measured from optical traces (ms) |
|---|---|---|---|---|
| 0.25 | 1094 | 2.218 | 5.48 | 5.3 ± 0.7 |
| 1 | 502.9 | 8.585 | 9.53 | 7.9 ± 2.1 |
| 2.25 | 343.2 | 17.69 | 13.33 | 10.7 ± 3.2 |
| 4 | 280.0 | 28.59 | 17.44 | 17.4 ± 3.1 |

### References:


1. K. Ohta, H. Ishida, Matrix formalism for calculation of the light beam intensity in stratified multilayered films, and its use in the analysis of emission spectra. *Appl. Opt.* **29**, 2466–2473 (1990).

2. J. Rivnay, *et al.*, Structural control of mixed ionic and electronic transport in conducting polymers. *Nature Communications* **7** (2016).

3. W. Shi, *et al.*, Micron-thick highly conductive PEDOT films synthesized via self-inhibited polymerization: roles of anions. *NPG Asia Materials* **9**, e405–e405 (2017).

4. D. A. Koutsouras, *et al.*, Impedance Spectroscopy of Spin-Cast and Electrochemically Deposited PEDOT:PSS Films on Microfabricated Electrodes with Various Areas. *ChemElectroChem* **4**, 2321–2327 (2017).

5. E. Stavrinidou, *et al.*, Direct measurement of ion mobility in a conducting polymer. *Adv. Mater.* **25**, 4488–4493 (2013).

6. H.-E. Tseng, T.-H. Jen, K.-Y. Peng, S.-A. Chen, Measurements of charge mobility and diffusion coefficient of conjugated electroluminescent polymers by time-of-flight method. *Applied Physics Letters* **84**, 1456–1458 (2004).

7. M. L. Seibenhener, M. W. Wooten, Isolation and culture of hippocampal neurons from prenatal mice. *J. Vis. Exp.* (2012) https:/doi.org/10.3791/3634.

8. J. B. Zuchero, Purification of dorsal root ganglion neurons from rat by immunopanning. *Cold Spring Harb. Protoc.* **2014**, 826–838 (2014).

9. J. B. Zuchero, Purification and culture of dorsal root ganglion neurons. *Cold Spring Harb. Protoc.* **2014**, 813–814 (2014).
